# DNA-scaffolded biomaterials enable modular and tunable control of cell-based cancer immunotherapies

**DOI:** 10.1101/587105

**Authors:** Xiao Huang, Jasper Z. Williams, Ryan Chang, Zhongbo Li, Eric Gai, David M. Patterson, Yu Wei, Wendell A. Lim, Tejal A. Desai

**Affiliations:** Department of Bioengineering and Therapeutic Sciences, University of California, San Francisco, San Francisco, California, USA; Department of Cellular and Molecular Pharmacology, University of California, San Francisco, San Francisco, California, USA; Department of Pharmaceutical Chemistry, University of California, San Francisco, San Francisco, California, USA

## Abstract

Advanced biomaterials provide versatile ways to spatially and temporally control immune cell activity, potentially enhancing their therapeutic potency and safety. Precise cell modulation demands multi-modal display of functional proteins with controlled densities on biomaterials. Here, we develop an artificial immune cell engager (AICE) platform – biodegradable particles onto which multiple proteins are densely loaded with ratiometric control via short nucleic acid tethers. We demonstrate the impact of AICE with varying ratios of anti-CD3 and anti-CD28 antibodies on *ex vivo* expansion of human primary T cells. We also show that AICE can be used to control the activity of engineered T cells *in vivo*. AICE injected intratumorally can provide a local priming signal for systemically administered AND-gate chimeric antigen receptor T cells, driving local tumor clearance while sparing uninjected tumors that model potentially cross-reactive healthy tissues. This modularly functionalized biomaterial thus provides a flexible platform to achieve sophisticated control over cell-based immunotherapies.

---

Functionalized biomaterials can work synergistically with natural or engineered immune cells for cancer immunotherapy^1–15^. Adoptive cell therapy (ACT) using chimeric antigen receptor (CAR) engineered T cells has shown success and clinical approval for the treatment of B cell cancers^16^. However, for CAR T cells to fulfill their promising potential, particularly for targeting solid tumors^17, 18^, important challenges must be overcome to improve both efficacy^17–20^ and safety^21–24^. For engineered anti-tumor T cells to achieve durable tumor remission in patients, a demanding manufacturing process is required to ensure the quantity and quality of the cell product for therapeutic use^1, 25–27^. To increase the tumor targeting specificity and avoid “on-target, off-tumor” toxicity in bystander healthy tissues^28^, CAR-T cells have been engineered with combinatorial antigen AND-gate activation control that requires sensing two antigens on a target cell to initiate killing. The clinical application of this strategy, however, requires systematic identification of tumor specific antigen combinations and would also benefit from better control of cell activity^27, 29^.

Biomaterials functionalized with modulatory biomolecules at pre-specified densities have been shown to communicate with and control therapeutic immune cells both *ex vivo* and *in* vivo^1–15^. For example, synthetic materials have been functionalized with agonistic antibodies for CD3 and CD28 to drive *ex vivo* T cell expansion^25, 26^. However, biodegradable materials^25^ with optimized material structure/composition and controlled surface moiety loading can provide unique opportunities to improve the ease of manufacturing and the quantity and quality of T cells produced. There is also a growing interest in the use of biomaterials for local modulation of engineered T cell activity *in vivo* during the course of treatment^1^.

Nonetheless, the use of biomaterials to precisely modulate immune cells still faces key challenges. There is a need for robust chemical conjugation strategies to surface-functionalize biodegradable materials with multiple signals (e.g. proteins/antibodies) at high densities and precisely controlled ratios^30–33^. PEG (polyethylene glycol) is commonly used as a linker or scaffold for the surface conjugation of biomolecules^31^, but this method is limited by the inefficient presentation of functional groups for attaching biomolecules due to PEG’s flexibility^34^. Synthetic short oligonucleotides - natural polymers with controllable sequence and structure - have been used as surface scaffolds on metallic particles for siRNA delivery^34–37^, but have yet to be fully utilized for modular payload assembly on biodegradable materials.

In this work, we developed short synthetic DNA scaffolds for the efficient and versatile functionalization of proteins or antibodies on the surface of biodegradable particles (**Fig. 1a**). A series of optimization studies were carried out to achieve maximum loading density, ratiometric control of moiety loading, adaptability to different particle size/composition, and feasibility for *in vivo* use. Micron-sized artificial immune cell engagers (AICE) were made from biocompatible poly(lactic-co-glycolic acid) (PLGA) polymer^38^ (**Fig. 1a**). We demonstrated that: **i**) we can load a range of immune modulators (e.g. anti-CD3, anti-CD28, and IL-2) on AICE at controlled ratios and densities; **ii**) the ratiometric control of costimulatory ligands is essential for optimal *ex vivo* T cell activation and expansion with higher yield and less exhaustion; **iii**) we can locally administer antigen-presenting AICE to control AND-gate CAR-T cell activation and tumor clearance *in vivo* (**Fig. 1a**). This modular materials-based strategy can provide versatile and precise synthetic control of natural or engineered immune cells for cancer immunotherapy^1, 12, 29^

**Figure 1.**
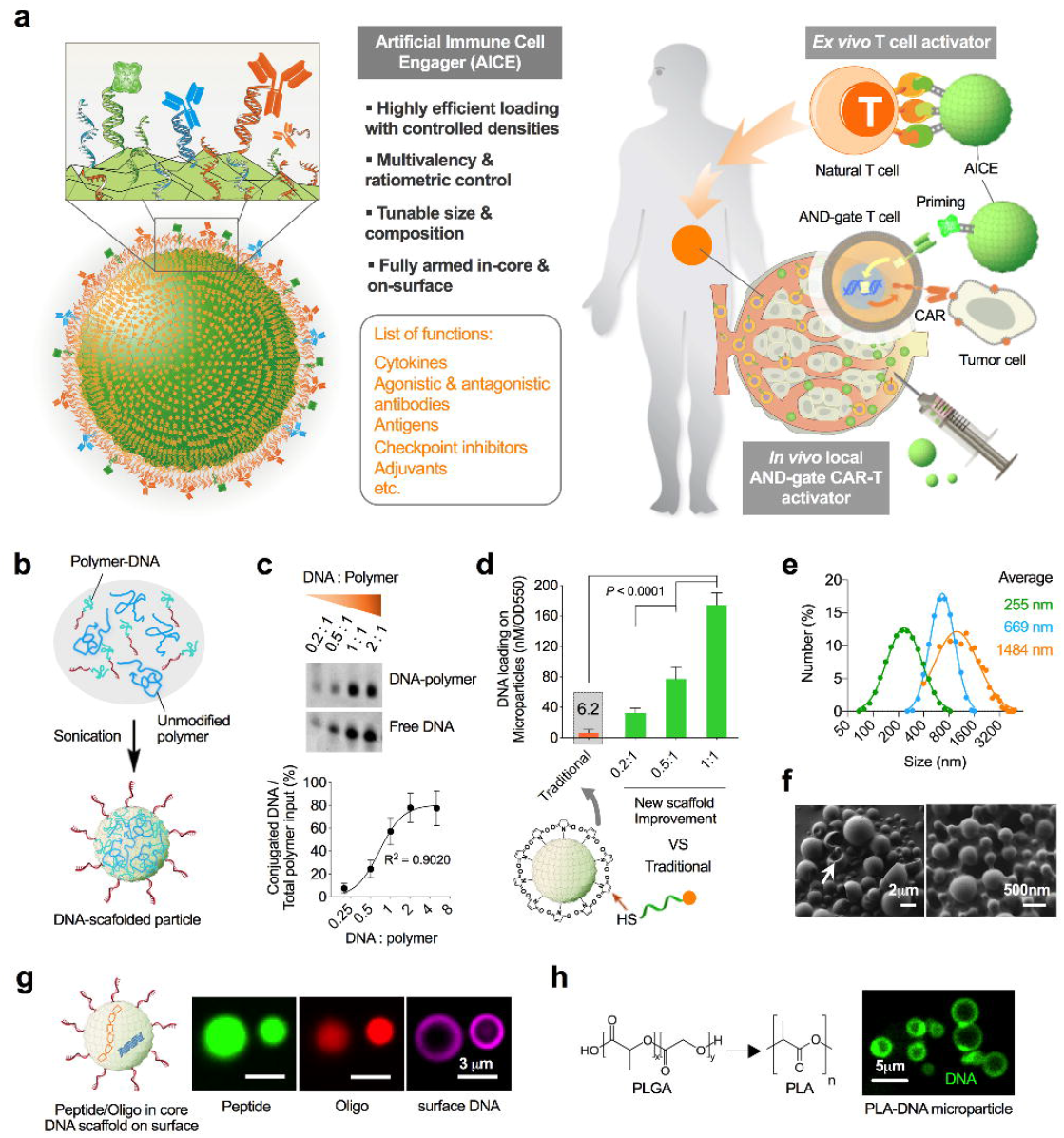
The synthesis of polymeric micro-/nano-particle with surface DNA scaffolds for therapeutic protein presentation. (**a**) Schematic of biodegradable polymeric particles with versatile therapeutic protein loading through surface DNA scaffolds, and their use as artificial immune cell engager (AICE) for *ex vivo* T cell expansion and *in vivo* activation control. (**b**) Diagram of DNA presentation on polymeric particles through the emulsion protocol. (**c**) Representative denatured PAGE electrophoresis gel image of the DNA-polymer conjugation reaction at different stoichiometry, and the quantification by densitometry analysis. Data are mean ± s.d. (n = 3 independent experiments), and the curve was fitted by Hill slope specific binding model. (**d**) Hybridized DNA scaffold on PLGA particles made from DNA-polymer conjugates of different coupling efficiencies, versus surface conjugation of dye-labeled DNA after particle fabrication. Data are mean ± s.d. (n = 6 independent samples from 3 independent experiments), and *P* values were determined by one-way analysis of variance (ANOVA) and Tukey’s tests. (**e**) Representative size distribution of DNA-decorated PLGA particles from different fabrication protocols described in Figure S1j (n = 3 independent experiments). Green and blue curve are through dynamic light scattering assay; orange curve is from ImageJ software analysis of confocal fluorescence images. (**f**) Representative scanning electron microscopy image of PLGA particles with average size at 1484 nm and 669 nm (n = 3 independent samples). White arrow points to the particle with concave shape at one side. (**g**) Representative images of PLGA microparticles with peptides and oligonucleotides in the core and DNA scaffold on the surface (n = 2 independent experiments). (**h**) Representative image of DNA-scaffolded microparticles composed of PLA polymer (n = 3 independent experiments).

## RESULTS

### Fabrication of surface DNA scaffolded polymeric particles

Synthetic short DNA strands were immobilized on the surface of polymeric particles to create an adaptable scaffold for loading bioactive molecules. The emulsion protocol of particle synthesis^38^ was modified to add conjugated polymer-DNA amphiphilic molecules during the fabrication process such that the hydrophilic DNA segment distributes around the hydrophobic core upon sonication (**Fig. 1b**). Polymer-DNA molecules were first synthesized through a conjugation of thiol-modified DNA to Maleimide (Mal)-modified polymer. We optimized the polymer, DNA length, solvent, and reaction conditions to generate optimal polymer-DNA molecules to form stable PLGA (50:50, MW 38-54k, Sigma) particles with dense DNA scaffolds (**Fig. Supplementary 1a-d**). PLGA10k-PEG5k-Mal was identified as an important polymer-DNA component to ultimately form microparticles of the desired size (1-2 μm in diameter), so we kept the polymer constant while varying the excess of DNA input during the conjugation reaction and observed an increase in polymer coupling efficiency (**Fig. 1c**).

The direct incorporation of polymer-DNA reaction mixtures with different DNA:polymer ratios into the particle fabrication protocol yielded a significant increase of surface payload-attachable DNA scaffold density, as determined by a fluorescence-based hybridization analysis (**Fig. 1d**). Strikingly, the highest average surface loading density on particles (~5 million DNA duplexes per particle, **Fig. Supplementary 1e-g**) was roughly analogous to the theoretical limit (at ~4 million by footprint calculation, based on ~2 nm diameter of DNA duplex) of a spherical particle with a 2 μm diameter. In particular, this hybridization-guided loading was about 27-fold more efficient than that from the traditional method^33^, in which thiol-modified DNA molecules were conjugated to surface-exposed Mal groups after particle fabrication using equal input amount of PLGA10k-PEG5k-Mal (**Fig. 1d**). The hybridization-guided biomolecule assembly protocol was optimized to a 30-minute incubation at 37°C (**Fig. Supplementary 1h**), and functionalized particles can be further lyophilized for storage and transportation (**Fig. Supplementary 1i**).

To accommodate diverse applications from intracellular payload delivery to extracellular signal transduction, DNA-scaffolded particles can be synthesized at various sizes and also loaded with biomolecules in their core. Particle size can be controlled by tuning key emulsion parameters (**Fig. 1e** and **Supplementary Fig. 1j**). We observed a concave shape on one side for some larger particles (~2 μm in diameter) but not for smaller particles (~500 nm) (**Fig. 1f**) possibly due to the high surface hydrophilicity with DNA scaffolds leading to higher surface to volume ratio. Additionally, we showed that the core of these DNA-scaffolded microparticles could also be loaded with biomolecules including oligonucleotides and peptides through a double-emulsion protocol (**Fig. 1g** and **Supplementary Fig. 1k**). This strategy of fabricating DNA-scaffolded particles was also successfully replicated using other polymers such as polylactic acid (PLA) (**Fig. 1h** and **Fig. Supplementary 1l,m**).

### Multi-functionalization of DNA-scaffolded particles

Biomolecules can be loaded on the surface of DNA-scaffolded particles using one of two strategies: a) linking functional groups of the scaffolds to biomolecules through the surface step-by-step conjugation via a bifunctional linker; and b) through the direct hybridization of complementary DNA-biomolecule conjugates to the scaffold (**Fig. 2a**). In the case of a fluorescently labeled human IgG, the surface conjugation strategy resulted in saturated protein loading even with increased densities of available DNA linkers on surface (**Fig 2b, Fig 1d** and **Fig. Supplementary 2a**). In contrast, the hybridization-guided assembly strategy showed a dramatically higher level of IgG loading for particles with denser DNA linkers, and the increase in IgG loading corresponded to the scaffold density (**Fig. 2b**). The highest level of IgG loading achieved (at ~0.6 million per particle) was again comparable to the theoretical footprint limit of a spherical particle at 2 μm in diameter (~0.64 million per particle, based on ~5 nm diameter of IgG) (**Fig. 2b**). For generating DNA-antibody conjugates with the minimum damage to the antibody activity, different chemistries (**Fig. Supplementary 2b,c**) were attempted and the “TCEP” strategy^6^ was identified as optimal, whereby the antibody hinge region was selectively reduced to expose the thiol group for 3’NH_2_-modified complimentary DNA to attach through a MAL-dPEG4-NHS linker (Quanta Biodesign). For example, anti-PD-L1 antibody conjugated with DNA through this strategy showed intact binding activity (**Fig. Supplementary 2c**), and its loading on PLGA microparticles resulted in high binding specificity for PD-L1 positive cells (**Fig. 2c**). For DNA tethering of proteins other than antibodies, other options are available to yield products with intact activity (**Fig. Supplementary 2d,e**).

**Figure 2.**
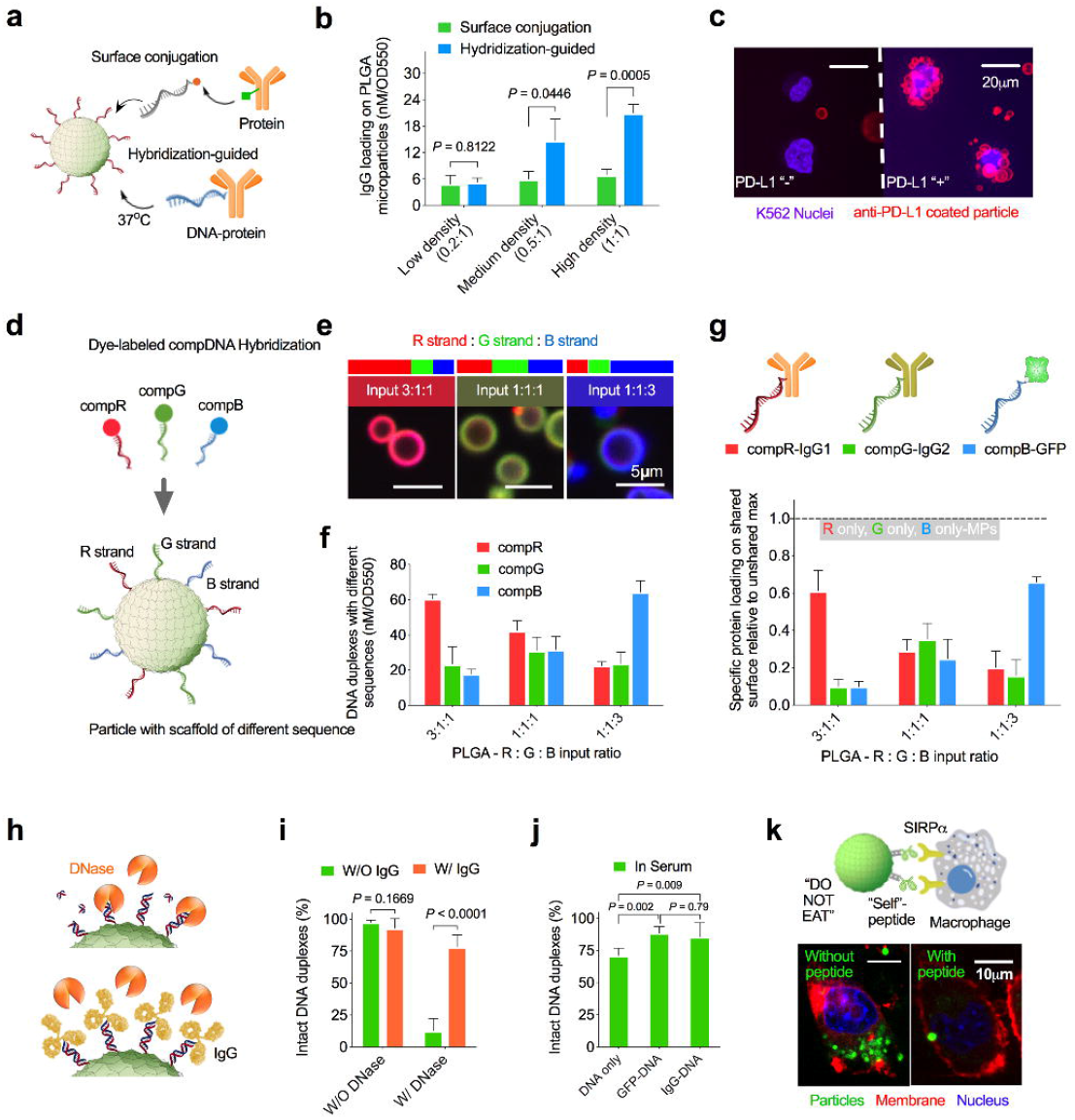
The multi-functionalization of DNA-scaffolded PLGA microparticles. (**a**) Schematic of two strategies of antibody loading on the particle surface. (**b**) Quantification of FITC-labeled IgG loading density by the two strategies depicted in (**a**). Data are mean ± s.d. (n = 3 independent experiments), and *P* values were determined by two-tailed paired *t* test. (**c**) Representative confocal microscopy images of anti-PD-L1 antibody loaded PLGA microparticles co-incubation with PD-L1 +/-K562 cells for 1 hour (n = 3 biologically independent samples per group). (**d**) Schematic of the control of DNA scaffolds with distinct sequences at intended ratios and its confirmation through corresponsive dye-labeled complementary DNA (compDNA) hybridization test. (**e**) Representative confocal microscopy images of PLGA microparticles with DNA scaffolds of different sequence compositions after hybridization with corresponding dye-labeled compDNA. The merged images of particles agreed well with the theoretically integrated color at different input ratios. (**f**) DNA duplexes of different sequences on particles by the fluorescence-based analysis. Data are mean ± s.d. (n = 5 independent samples from 2 independent experiments). (**g**) Loading density of three different proteins sharing the same surface guided by DNA scaffold compositions, relative to their individual maximum without sharing. Data are mean ± s.d. (n = 3 independent experiments). (**h**) Schematic of DNA scaffold integrity on the particle surface with and without IgG coverage, in the presence of DNase. (**i**) Intact DNA duplexes on PLGA microparticles after co-incubating with large excess of DNase for 20 minutes at 37°C. Data are mean ± s.d. (n = 3 independent experiments), and *P* values were determined by two-tailed paired *t* test. (**j**) Intact DNA duplexes with different attachments on PLGA microparticles after incubation in human serum for 1 hour at 37°C. Data are mean ± s.d. (n = 5 independent samples from 2 independent experiments) and *P* values were determined by one-way ANOVA and Tukey’s tests. (**k**) Representative confocal microscope images of macrophages treated with particles with and without “Self’-peptide presentation (n = 3 biologically independent samples per group).

Another advantageous feature of using DNA as a surface scaffold is that the uniqueness of nucleotide sequences enables the independent control of loading multiple cargos on the same material surface (**Fig 2d**). Polymer-DNA conjugates with three distinct DNA sequences (namely R, G, and B strand) that do not anneal with each other (**Fig. Supplementary 2f**) were synthesized and incorporated into particle emulsion with reaction mixtures at different ratios (3:1:1, 1:1:1 and 1:1:3). After surface hybridization with a mixture of their respective dye-labeled complementary strands (**Fig. 2d**), we found that the ratios of hybridized DNA scaffolds of different sequences on PLGA microparticles were consistent with the polymer-DNA conjugate input, evidenced by both confocal fluorescence imaging (**Fig. 2e**) and a fluorescence-based quantification assay (**Fig. 2f**). The ratiometric control of dye-labeled DNA strand loading was equally efficient for PLA microparticles (**Fig. Supplementary 2g,h**). Based on this, we co-loaded three different proteins (GFP and two antibodies), each individually attached with one of the three complementary DNA strands, onto PLGA microparticles with varying ratios of the DNA surface scaffold sequences above. Similar distribution of each protein cargo compared with the scaffold population was observed (**Fig. 2g**), demonstrating the robust ratiometric control of surface protein loading using this particle synthesis strategy.

To enable *in vivo* use, we explored the stability of the DNA scaffolds on the surface of PLGA microparticles in the presence of a large excess of DNase. (**Fig. 2h**). We observed an almost complete (~87%) degradation of DNA scaffolds after DNase treatment; however, this dropped down to ~20% with IgG molecules conjugated to the scaffolds through the surface step-by-step conjugation (~1/4 of the highest IgG loading through hybridization-guided assembly) (**Fig. 2b,i**). This suggests that the surface-loaded proteins are able to protect the DNA scaffolds. In human serum, we did not observe as significant DNA degradation for the naked particles due to the lower DNase concentration in serum than used in **Fig.2i**; however, the degree of DNA degradation similarly decreased as surface payload proteins were attached (**Fig. 2j**).

Macrophage uptake is a major obstacle^39^ for particle-based drug delivery *in vivo*, therefore we incorporated a CD47-mimic “self”-peptide^40^ on PLGA microparticles (**Fig. 2k**). Fluorescein-labeled PLGA microparticles with and without surface-loaded “self”-peptide were co-incubated with a mouse macrophage line J774A.1 *in vitro*. The particle uptake assay showed that the “self”-peptide decoration significantly reduced the uptake of particles by macrophages (**Fig. 2k** and **Supplementary Fig 2i,j**). This “self”-peptide functionalization provides a possible strategy to reduce the clearance of particles *in vivo*.

### Design of particles with ratiometrically controlled moieties for *ex vivo* human T cell activation

Given the ability to synthesize particles with ratiometric control of multiple surface cargos, we loaded anti-CD3 and anti-CD28 antibodies on AICE (PLGA microparticles) at varying ratios from 1:5 to 5:1. Anti-CD3/CD28 AICE were compared to commercial anti-CD3/CD28 Dynabeads at the same particle to cell ratio for the ability to expand human primary T cells with minimized differentiation and exhaustion (**Fig. 3a**). Functionalized AICE activated T cells (**Fig. 3b**) and yielded higher or equivalent T cell expansion compared to Dynabeads expansion across three human T cell donors (**Fig. 3c,d**). Although there were large donor-to-donor differences (also observed for Dynabeads), we observed a linear increase of cell yield from AICE-[1:5, anti-CD3:anti-CD28] to AICE-[3:1] for both CD4+ and CD8+ cells at day 14 (**Fig. 3c** and **Fig. Supplementary 3a**). The phenotype of expanded T cells at day 14 was then explored by measuring expression of CD45RA and CCR7 surface markers (**Fig. 3e**). Interestingly, the population distribution of the 4 differentiation states (naïve, central memory [CM], effector memory [EM] and effector memory RA [EMRA]), displayed a pattern (**Fig. 3f** and **Fig. Supplementary 3b**) corresponding to the cell expansion trend among AICE with different anti-CD3 to anti-CD28 ratios (**Fig. 3c**). Through staining for T cell exhaustion markers LAG-3, PD-1 and TIM-3, we found that the population of exhausted cells at the optimal condition AICE-[3:1] was generally less than those activated by Dynabeads among the three donors (**Fig. Supplementary 3c,d**). Taken together, the data demonstrates that the surface ratio control of functional moieties on synthetic materials is important for the quantity and phenotype of cells yielded from *ex vivo* T cell expansion.

**Figure 3.**
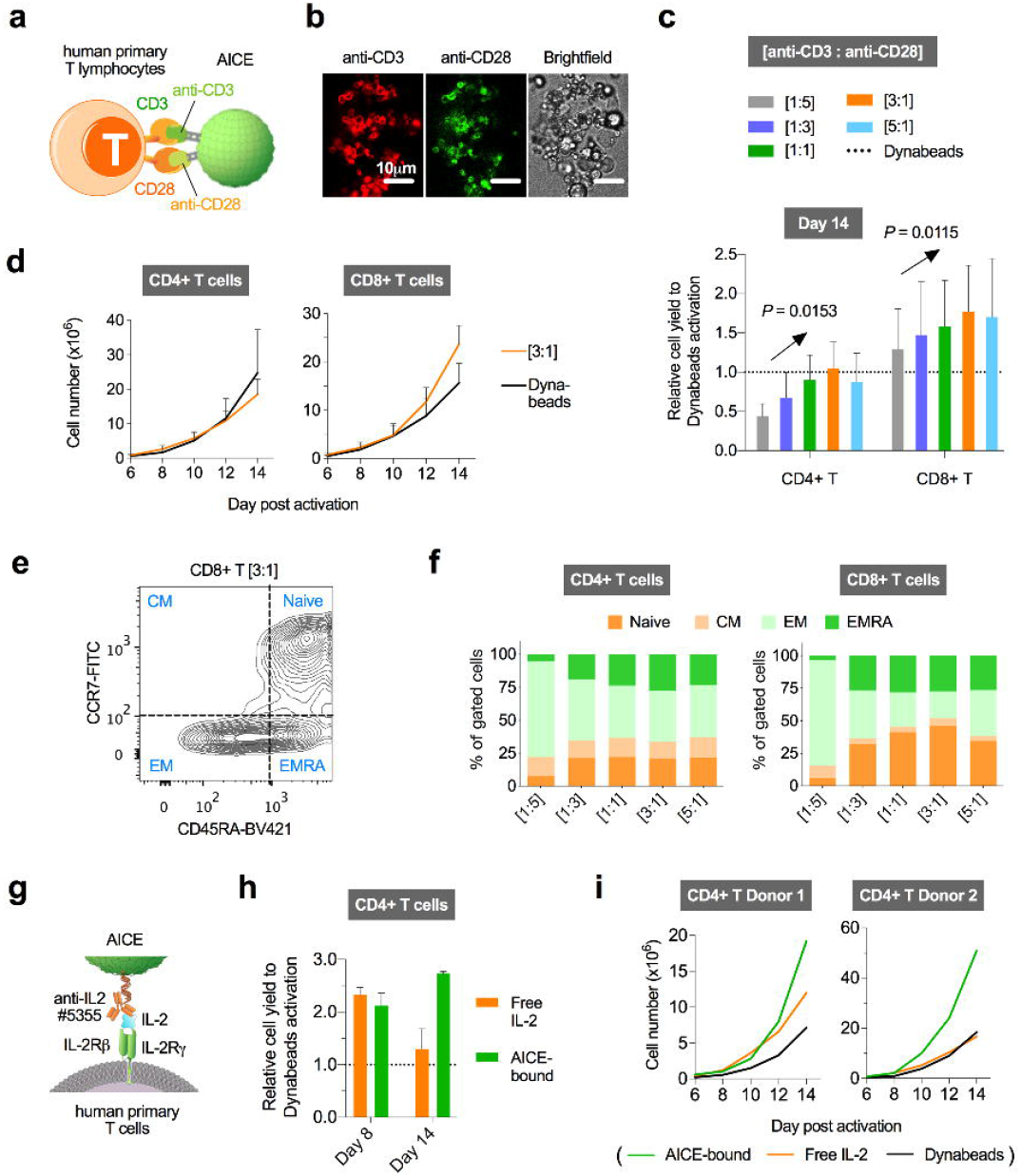
AICE with ratiometrically controlled moieties for human primary T lymphocytes *ex vivo* expansion. (**a**) Schematic of AICE (in this figure PLGA microparticles loaded with anti-CD3 and anti-CD28 antibodies of varying ratios from [1:5] to [5:1]) for primary T cell expansion, compared with commercial available T cell expander Dynabeads (Invitrogen). (**b**) Representative confocal microscopy images of human primary CD8+ T cell co-incubated with AICE [1:1] for overnight, showing AICE-induced cell clumps (n = 3 biologically independent samples). (c) Cell yield of CD4+ and CD8+ T cells at day 14 after activation by AICE with anti-CD3 and anti-CD28 antibodies at different ratios in (**a**), relative to Dynabeads. Data are mean ± s.e.m., and *P* values were determined by one-way ANOVA test for linear trend (n = 3 independent donors of two independent experiments). (**d**) Growth curve of primary T cells from three different donors upon the activation by AICE [3:1], compared to Dynabeads control. Data are mean ± s.e.m. (n = 3 independent donors of two independent experiments). (**e**) Gating strategy through CCR7 and CD45RA expression levels for distinguishing cells at specific differentiation stages: naïve, central memory (CM), effector memory (EM), and effector memory RA (EMRA). (**f**) Differentiation profile of T cells 14 days after activation by AICE of varying ratios of surface moieties. Data are mean of n = 3 independent donors. (**g**) Schematic of IL-2 presented on particles through its antibody (clone5355) that exposes the epitope for β and γ units of its receptor on T cells, promoting the cell proliferation. (**h**) Cell yield of primary CD4+ T cells activated by AICE [3:1] with surface-bound IL-2 or free IL-2 at day 8 and 14, relative to the Dynabeads method. Data are mean ± s.d. (n = 2 independent donors). (i) Growth curve of CD4+ primary T cells from two different donors treated with AICE [3:1] together with surface bound IL-2 or free IL-2, compared to Dynabeads control. Data are mean of n = 2 technical replicates.

As an alternative to the standard protocol of supplementing free IL-2 in the media for *ex vivo* T cell culture^25^, we loaded IL-2 on AICE through the surface presentation via its antibody clone #5355 (**Fig. Supplementary 3e**). This particular anti-IL-2 antibody was engineered to facilitate the binding of IL-2 to its β and γ receptor on T cells thus promoting the proliferation of non-Treg T cells^41^ (**Fig. 3g** and **Fig. Supplementary 3f**). Using the optimal condition of AICE[3:1] (anti-CD3:anti-CD28) determined above, we compared primary T cell expansion by supplementing equal amount of free IL-2 versus surface bound IL-2 (**Fig. Supplementary 3e**). Surface IL-2 loading enhanced CD4+ and CD8+ T cell expansion, and particularly improved expansion of CD4+ primary T cells after day 8 (**Fig. 3h,i** and **Fig. Supplementary 3a,g**). Although with higher level of cell yield, the populations of LAG-3 and PD-1 positive CD4+ T cells still decreased in the two tested donors (**Fig. Supplementary 3h**), demonstrating the advantages of AICE-surface presentation of IL-2 for *ex vivo* expansion.

### Design of particles to prime activation of AND-gate CAR-T cells

We also explored the use of AICE coated with an orthogonal antigen (GFP) to prime AND-gate CAR-T cell tumor recognition circuits, with the goal of preventing “on-target off-tumor” toxicity by restricting cell killing to tumors locally-injected with AICE microparticles (**Fig 4a**). These AND-gate T cells utilize a modular synthetic Notch (synNotch) receptor with an extracellular domain to recognize a target antigen, and an intracellular transcriptional activator (TF) domain to control expression of a CAR targeting a second antigen^28^. Killing is only induced when both antigens are presented to T cells, with one activating the synNotch receptor to release the TF domain for CAR expression and the other activating CAR-mediated cytotoxicity^28^. Herein, human primary T cells were transduced with a gene encoding a constitutively-expressed synNotch receptor with an extracellular anti-GFP nanobody and an intracellular Gal4DBD-VP64 synthetic transcription factor (TF) domain, together with a TF-inducible CAR gene targeting HER2 (human epithelial growth factor receptor 2)^28^ (**Fig. 4a**). In this system, T cells should only express the anti-HER2 CAR and kill HER2 positive target cells after priming via the anti-GFP synNotch binding AICE-presented GFP.

**Figure 4.**
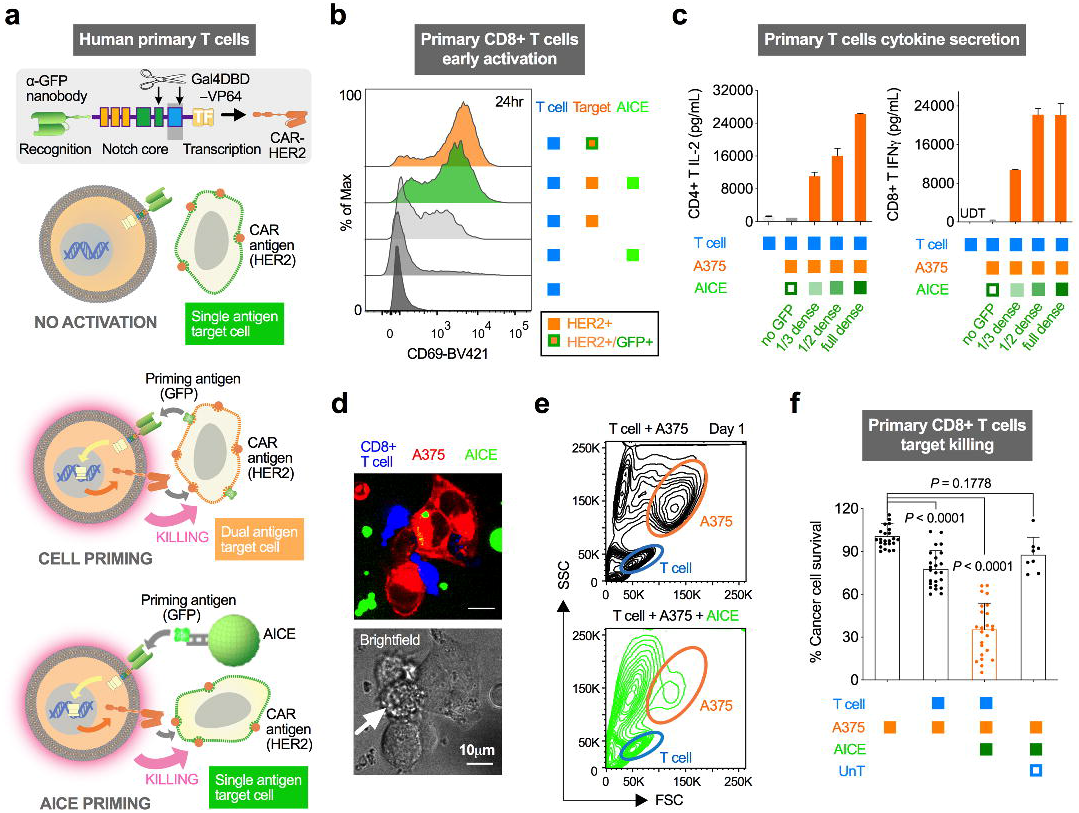
Activation of AND-gate engineered human primary T cells through AICE functionalized with priming antigens for the induction of CAR toxicity. (**a**) Schematic of T cell engineering for dual antigen AND-gate targeting, and its priming by synthetic antigen (herein GFP) presented by AICE to trigger HER2-CAR guided killing, alternative to existing strategy of dual antigen target cell killing. (**b**) Expression level of CD69 as an early activation marker for CD8+ synNotch/CAR T cells after 24 hours of co-incubation with AICE and HER2-overexpressed A375 target cells, compared to the treatment of target cell K562 with GFP and HER2 co-expressed (n = 2 independent experiments). (**c**) IL-2 and IFN-γ secretion levels by primary CD4+ and CD8+ T cells 48 hours after AICE and HER2-overexpressed target cell A375 co-incubation. Data are mean ± s.d. (n = 2 technical replicates from 4 biologically independent samples). (**d**) Representative confocal fluorescence microscopy image of the co-incubation of CD8+ synNotch CAR-T cells, GFP-presenting AICE and HER2-overexpressing A375 cells after 24 hours (n = 2 independent experiments). The arrow points to a dead A375 cell. (**e**) Representative forward and side scatter flow cytometry plots after 24-hour co-culture of HER2-expressing A375 and CD8+ synNotch CAR-T cells in the presence and absence of AICE (n = 3 independent experiments). The T cells fall within the blue gate, and the target A375 cells are in the orange gate. (**f**) Cell survival test of target A375 cells 48 hours after co-culture of AICE and synNotch CAR-T cells. Data are mean ± s.d. (n = 6 independent experiment with 4 independent T cell donors), and *P* values were determined by one-way ANOVA and Tukey’s tests.

Human primary CD8+ T cells engineered with the above circuit were co-cultured for 24 hours with single antigen target cells (human melanoma cell line A375 with HER2 overexpressed) in the presence or absence of GFP+ AICE microparticles. SynNotch/CAR T cells showed selective elevation of the early activation marker CD69 in the presence of both HER2+ A375 cells and GFP+ AICE, and the activation was similar to T cells exposed to dual antigen target cells (human leukemia cell line K562 with GFP and HER2 co-expressed) (**Fig. 4b**). For both CD4+ and CD8+ engineered primary T cells, we observed selective secretion of cytokines (IL-2 and IFN-γ, respectively) after 48 hours of co-culture with both HER2+ A375 cells and GFP+ AICE (**Fig. 4c**). Increasing AICE GFP density led to elevated levels of cytokine secretion (**Fig. 4c, Fig. Supplementary 4a**). Noticeably, T cells secreted significantly more cytokines with AICE presenting synNotch-priming GFP than with target cells presenting both synNotch and CAR antigens (A375 with GFP and HER2 co-expressed) (**Fig. Supplementary 4b**).

Human primary CD8+ T cells containing the same circuit as above also showed AND-gate killing behavior, selectively killing HER2+ A375 cells in the presence of GFP+ AICE microparticles. Co-incubation of GFP+ AICE and CD8+ human primary T cells led to some killing of co-cultured HER2+ A375 cells (with 1:1 ratio) at 24 hours (**Fig. 4d,e**), and the killing became more apparent after 48 hours with higher densities of priming signals on AICE (**Fig. 4f and Fig. Supplementary 4c-f**). In contrast, initial attempts using particles synthesized by traditional conjugation chemistry and with far lower GFP density failed to activate AND-gate CAR-T cells, likely due to the inadequate ligand density for robust synNotch activation. As a demonstration of adaptability for translational use, a synthetic short peptide (PNE peptide) computationally predicted to be non-immunogenic^42^ was engineered on AICE and co-incubated with its corresponding anti-PNE synNotch/anti-HER2 CAR CD8+ T cells and HER2+ target cells, and killing was significantly enhanced by PNE+ AICE addition (**Fig. Supplementary 4g**). The co-incubation of target cells with synNotch CAR-T cells in the absence of AICE also showed moderate cytotoxicity (**Fig. 4f and Fig. Supplementary 4c-g**), which is likely due to the leakiness of inducible CAR expression also observed from previous reports and can be improved through further optimization of synNotch receptors.

### Local tumor eradication in mouse model by the combinatorial treatment of priming particles and AND-gate T cells

To determine the feasibility of using local particle injections to send spatially-defined signals to therapeutic cells *in vivo*, we first monitored the distribution of intratumorally-injected AICE microparticles. We labeled surface DNA and polymer core with two distinct infrared dyes (Quasar705 and IR800CW respectively, **Fig. Supplementary 5a**) on microparticles composed of PLGA (~1.5 μm in averaged diameter) and PLA polymer (~1.9 μm in averaged diameter). AICE particle retention and half-life were then explored *in vivo* by injection into subcutaneous K562 xenograft tumor in NSG mice followed by the IVIS near-infrared fluorescence imaging at different time points. AICE were retained locally within the tumor for over a week (**Fig. 5a and Fig. Supplementary 5b**), and the fluorescence quantification showed an approximate half-life of 4-5 days for the surface coating of both types of particles (**Fig. 5b and Fig. Supplementary 5c-e**). These results demonstrate that AICE injection can be used to send localized, sustained signals to therapeutic cells during the course of treatment.

**Figure 5.**
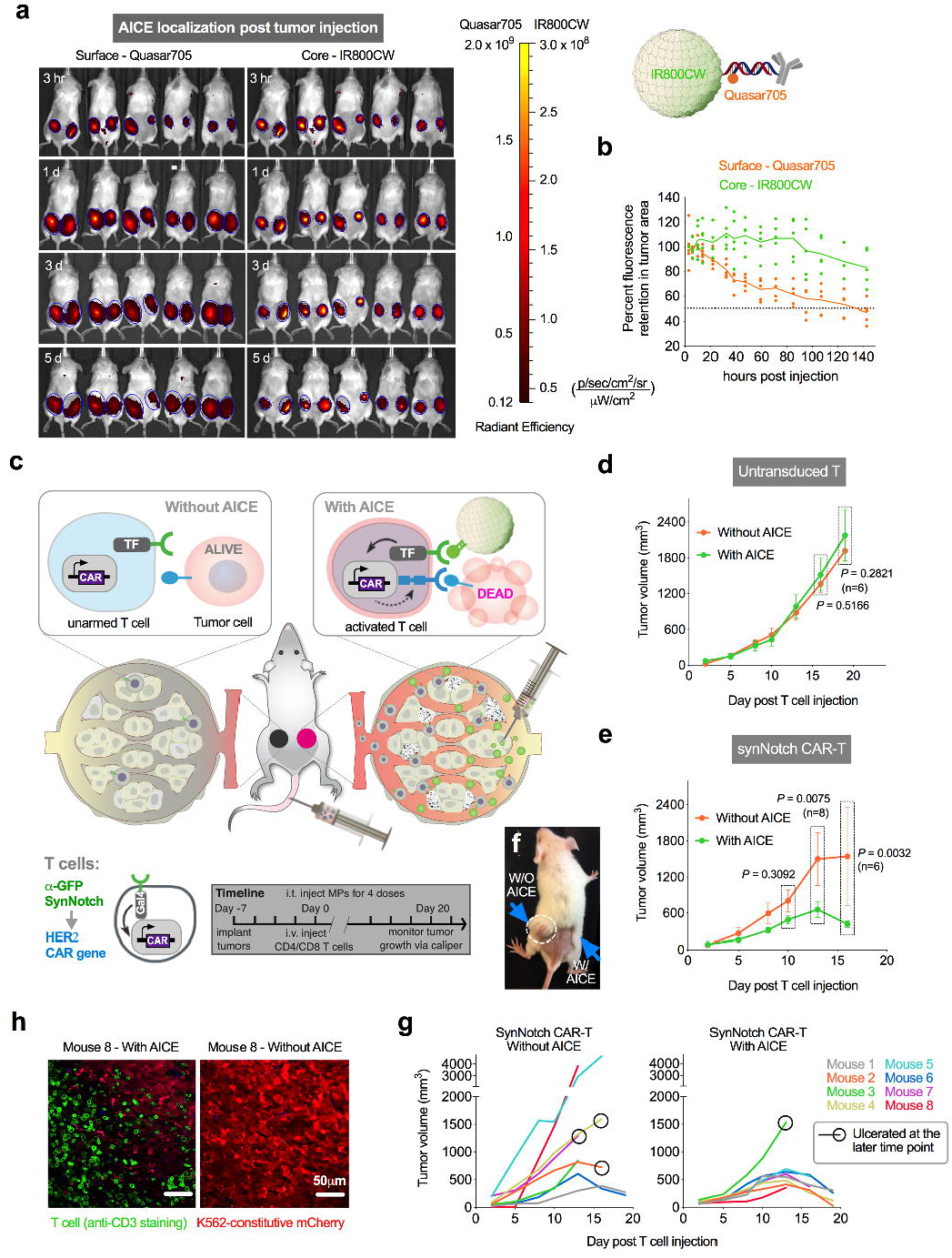
Selective tumor killing *in vivo* by local activation of synNotch CAR-T cells using AICE. (**a**) IVIS fluorescence imaging of IR dye-labeled AICE (Quasar705 on surface DNA and IR800CW in core) in subcutaneous K562 tumors of NSG mice over 6 days (n = 5 mice). Left tumors are injected with AICE made of PLA; right tumors are injected with AICE made of PLGA. (**b**) Quantification of the average signal intensity at the tumor area (right tumor with AICE-PLGA) for the full set IVIS images acquired over 6 days in (**a**). Lines are the mean signal from n = 5 mice. (**c**) Schematic of two tumor model for selected clearance by AICE-primed synNotch CAR-T cell activation through local intratumoral injection, and the overall treatment timeline. (**d**) Tumor volume of AICE-injected tumor versus contralateral tumor over 19 days after combinatorial treatment of untransduced primary T cell and GFP-coated AICE. Data are mean ± s.e.m. (n = 6 mice), and *P* values were determined by two-tailed paired *t* test. (**e**) Tumor volume of AICE-injected tumor versus the contralateral tumor over 19 days after combinatorial treatment of synNotch CAR-T cell and GFP-coated AICE. Data are mean ± s.e.m. (n = 8 mice, and 2 mice met the euthanasia criteria at day 13-15), and *P* values were determined by two-tailed paired *t* test. (**f**) Representative image of the mice intravenously injected with synNotch CAR-T cells, showing diminished tumor on AICE injection site but growing tumor at the contralateral flank. (**g**) Tumor volume of individual mice shown in (**e**). (**h**) Representative images of fixed staining of CD3+ T cells in tumors collected from mouse 8 in (**g**), which is sacrificed in earlier date meeting euthanasia criteria (n = 10 technical replicates).

We hypothesized that AICE made of GFP decorated PLGA microparticles could be injected intratumorally as a local activator for systemically administered anti-GFP synNotch/anti-HER2 CAR T cells (**Fig. 5c**). NSG mice were implanted subcutaneously with the same HER2-overexpressed K562 xenograft tumors in bilateral flanks as a model for a tumor and healthy tissue both expressing the CAR antigen. After a week, mice were administered synNotch CAR-T cells once intravenously in combination with AICE intratumorally injected for 4 doses (**Fig. 5c**). The size of AICE-injected ipsilateral tumors decreased over time, in contrast to the contralateral tumors within the same mice without AICE injection and tumors in mice injected with AICE plus untransduced primary T cells (**Fig. 5d-g**). We also performed fluorescence microscopy on fixed tumor samples of mice sacrificed at an early timepoint and observed selective T cell infiltration in the AICE-injected tumor (**Fig. 5h and Supplementary 5f**). These results demonstrate that AICE can provide a spatially controlled signal *in vivo* for the local activation of synNotch CAR-T cells and induction of precision tumor clearance with limited risk of cross-reaction against healthy tissues.

## DISCUSSION

In this work, we demonstrate a unique platform using synthetic short DNA oligonucleotides as surface scaffolds on biodegradable materials for the precise and controlled loading of multiple biomolecules at specific ratio and density. These advanced biomaterials possess the ability to regulate immune cells in the context of *ex vivo* and *in vivo* activation of both natural and engineered T cells. Through this highly modular platform, biocompatible materials can be employed as substrates for a wide spectrum of modulatory biomolecules, including cytokines^5^, antigens^10^, checkpoint inhibitors^43^, agonistic or antagonistic antibodies^44^, adjuvants^9, 11^, etc., to regulate the local environment and enhance the efficacy of immune cells in cancer immunotherapy^8^. Synthetic materials with tunable properties can enhance the pharmacokinetics of regulatory biomolecules *in vivo*^1, 6, 8^, and sometimes provide a necessary surface substrate for biomolecules to activate the cognate cell signaling^10, 25, 28^. Biomaterials fabricated with controlled size, shape, and composition - for example, microspheres, nanowires, and porous scaffolds - can also serve as a local adjuvant^2, 3, 7, 13^ for biomolecules or engineered immune cells to exert tumor targeted toxicity without systemic diffusion, thus minimizing the off-target side effects. This can make a particular difference for solid tumor treatment, where functionalized biomaterials could be administered locally or engrafted post-surgery^3, 17, 45^.

Here we explored using a natural polymer, DNA, as the surface scaffold for loading biomolecules onto particles. We demonstrated, through near surface-saturated biomolecule loading and precise ratiometric control of moiety loading, that hybridization-based thermodynamics can overcome limitations of traditional surface conjugation such as reduced efficiency of multi-step reactions, steric hindrance, and decay of functional groups. The increased loading density of surface biomolecules was found to be necessary to make particles capable of presenting antigens to robustly activate synNotch receptors in primary human T cells, and may also significantly enhance immune cell signaling where ligand clustering is required^10, 46^. We also show the versatility of this strategy by fabricating particles of different sizes, as well as compositions with different degradation profiles. This method can be applied to other formats of biomaterial functionalization (e.g. scaffolds and nonspherical particles) where fragile biomolecules need to be presented at a material interface.

The densely-packed surface DNA scaffolds and the payload attachments protected DNA linkers from enzymatic degradation, which shows great promise for *in vivo* use. This organized packing of the particle surface with a protein corona may also minimize interactions with serum proteins to prevent macrophage clearance^47^. Furthermore, particle uptake by macrophages can be attenuated through the immobilization of a CD47 mimic “self”-peptide^40^. Safety concerns about immunogenicity from the DNA scaffolds can be addressed through optimization of sequences and special modifications^48^.

The precise ratio of agonistic antibodies (anti-CD3 and anti-CD28) for T cell activation on AICE impacts *ex vivo* T expansion efficiency, as well as T cell differentiation and exhaustion phenotypes. The ability to accurately control the ratio of multi-cargo particle loading should enable identification of novel, precise combinations of modulatory signals that lead to diversified and improved T cell behavior. The use of biodegradable materials also omits the need for synthetic material removal prior to cell implantation. Cumulatively, these properties suggest that AICE could aid in manufacturing for ACT by improving the ease-of-production and fine-tuning the yield and phenotype of the therapeutic cell product.

We and other groups have developed strategies to engineer T cells with multiple receptors to detect combinations of antigens^27–29^. All prior combinatorial antigen recognition systems have targeted combinations of antigens expressed on tumor cells and/or tumor-related cells^27, 29^. Here, we demonstrate a method to engineer T cells to recognize combinations of endogenous and orthogonal antigens presented by tumor cells and biocompatible materials, respectively. By incorporating material-presented antigen into combinatorial recognition circuits, we enable real-time controlled local activation of these circuits that can be achieved through dynamic particle dosing in living organisms/patients. Local injection of our particles serves to safely control receptor activation and downstream behavior of therapeutic cells *in vivo*. We demonstrated that synNotch-based combinatorial antigen recognition circuits can incorporate material-presented antigen sensing, and it is likely that other combinatorial antigen recognition systems, for example, chimeric costimulatory receptors (CCRs)^49^ in AND-gates or inhibitory CARs (iCARs)^50^ in NOT-gates, can as well.

We believe that the true promise of the AICE platform will be realized by using particles to present multiple signals within the tumor microenvironment. AICE particles could kick-start tumor killing by triggering CAR expression by synNotch-CAR T cells. T cell activity within the tumor could be further augmented by co-functionalization of AICE particles with additional cargos that go beyond a synNotch ligand, for instance, a costimulatory ligand and a checkpoint inhibitor tailored to circumvent the specific immune evasion mechanisms of a particular tumor type^17, 45^. Ultimately, it would be ideal to utilize intratumoral injection of AICE particles to initiate antigen-specific local immunity that could be converted into antigen spreading and systemic immunity by incorporating other cues onto/into the particles to stimulate innate immunity^18^.

## METHODS

Methods, including statements of data availability and any associated accession codes and references, are available in the online version of the paper.

### Note

*Any Supplementary Information and Source Data files are available in the online version of the paper*.

## ACKNOWLEDGMENTS

We thank Z. Gartner for DNA synthesis, S. Habelitz for DLS analysis, C. Hayzelden for SEM imaging, B. Hann for IVIS imaging, and C. Zamecnik and A. Li for helpful discussion. X.H. was supported by a UCSF program for breakthrough biomedical research (PBBR) postdoc independent research grant and a Li foundation fellowship. J.Z.W. was supported by a Genentech Pre-Doctoral Fellowship. R.C. was supported by National Institute of General Medical Sciences (NIGMS) Medical Scientist Training Program #T32GM007618.

## AUTHOR CONTRIBUTIONS

X.H., J.Z.W., R.C., W.A.L. and T.A.D. designed the experiments and interpreted the results. X.H., J.Z.W., R.C., Z.L., E.G., and Y.W. performed the experiments, and D.M.P. contributed to material designs and synthesis. X.H. analyzed the data and drafted the manuscript. X.H., J.Z.W., R.C., W.A.L. and T.A.D. edited the manuscript.

## ONLINE METHODS

### Synthesis of Thiol-modified DNA

3’Thiol-modified DNA were synthesized on 3’thiol-modifier 6 S-S CPG beads (Glen Research #10-1936-02) using an Expedite 8909 DNA synthesizer. DNA oligos were retrieved from the beads by 70°C incubation for 20 mins in the presence of AMA solution (Ammonium hydroxide:Methylamine=1:1,v/v), followed by vacuum evaporation (SpeedVac, ThermoFisher #SPD121P-230) for 3 hours to remove AMA. DNA oligos were reconstituted in TE buffer (Tris-EDTA, 10mM, pH7.5) and then filtered through 0.22 μm filter (Millipore #UFC30GV00), and stored at −20 °C.

### Synthesis of polymer-DNA amphiphilic molecule

3’Thiol-modified DNA oligos were deprotected by treating with 100x molar excess of Tris(2-carboxyethyl)phosphine hydrochloride solution (TCEP, Sigma #646547) at 37 °C for 1 hour, and purified by size-exclusion chromatography (Glen Research #61-5010). Unless otherwise specified, freshly prepared thiol-DNA 17mer at 200 μM were reacted with poly(lactide-co-glycolide)-b-poly(ethylene glycol)-maleimide (Mw 10,000:5000 Da, PLGA10k-PEG5k-Mal, Akina #AI053) at 1:1 ratio in the solvent of dimethylformamide (DMF)/H_2_O (90:10,v/v) with the addition of 0.2% triethanolamine, and incubated at room temperature overnight. The next day the solvent was removed by vacuum evaporation at 70°C for 3 hours, and the dried mixture was stored at −20°C. The product was verified by TBE-Urea gel electrophoresis (15%, ThermoFisher #EC68855) with SYBR gold dye staining (ThermoFisher #S11494), and the conjugation efficiency was quantified by gel densitometry analysis.

### Fabrication of polymeric particles with DNA scaffolds

For the fabrication of PLGA particles with varying sizes (**Fig. Supplementary 1j**), dried polymer-DNA reaction mixture (from 100 nmol of each reactant) reconstituted in 200 μL solvent (ethyl acetate:H_2_O=1:1,v/v) was incorporated into the mixture of 0.5 mL ethyl acetate (EtOAc) with certain amount of unmodified polymer (Poly-lactic-co-glycolic acid, 50:50, Mw 38,000-54,000, Sigma #719900), and 0.5 or 1.0 mL aqueous buffer (10mM sodium citrate, 600 mM Na+, pH3.0) with or without 1% polyvinyl alcohol (PVA, Sigma #81381). The whole mixture was then vortexed and probe-sonicated (Fisher miniRoto S56) on ice at 7-8 W for 5 × 5 s with 10 s intervals, and immediately added with 9 mL 0.2% PVA and stirred in the hood for 3 hours for ethyl acetate to evaporate. Particles were filtered through 40 μm cell strainer (Sigma #CLS431750), and washed through the centrifugation at 10,000x g for 10 mins followed by the resuspension of pellet in TE buffer (10mM Tris-HCl, pH8.0) with 0.01% Tween-20 added. This washing protocol stayed the same to later fabrications. After three washes, particles were suspended in TE buffer with 1% PVA and lyophilized (VirTis Advantage Plus) for long term storage. Particle size profile was characterized by dynamic light scattering (DLS) with a Malvern Nano ZS instrument, and SEM imaging using a Zeiss Ultra-55 scanning electron microscope. Particle concentration was normalized by the optical density at 550 nm (OD550) measured from a UV-Vis spectrometer (Molecular Devices SpectraMax M5).

PLA microparticles fabrication used the same protocol but adjusted emulsion mixture: 0.5 mL dichloromethane (DCM) with 100 mg/mL polylactic acid (Mw 60,000, Sigma #38534), 0.5 mL aqueous buffer (10 mM sodium citrate, 600 mM Na+, pH3.0) and the polymer-DNA reaction (from 200 nmol of each reactant) reconstituted in 200 μL solvent (ethyl acetate:H_2_O=1:1,v/v). To encapsulate biomolecules in the core of DNA-scaffolded particles, another emulsion step was added prior to the above emulsion: biomolecules including 0.25 mg FITC-labeled peptide with 21 amino acids (LifeTein) and 50 nmol 5’Cy3-labeled DNA-21mer (Bioresearch Technologies) dissolved in 50 μL PBS (phosphate buffered saline, pH7.0) was mixed into 0.5 mL ethyl acetate with dissolved PLGA (Sigma #719900), and probe-sonicated at 7-8 W for 5 × 5 s with 10 s intervals on ice. Immediately after this, amphiphilic polymer-DNA and aqueous buffer was added following the same protocol described above.

### Attachment of complementary DNA to proteins and their purification

Antibodies were attached to complementary DNA strands through the “TCEP” strategy. Various modified complementary DNA strands were all from Biosearch Technologies. Antibodies were first exchanged in reducing buffer (PBS with 10 mM EDTA) by size exclusion chromatography (Zeba Spin Desalting Column, ThermoFisher #89882), and the disulfide bond at the hinge region was selectively reduced by adding TCEP with 3 molar or 4.5 molar excess and incubating at 37°C for 1 hour. Excess TCEP was removed through size exclusion chromatography. 3’-NH2 modified complementary DNA was conjugated with a MAL-dPEG4-NHS linker (Quanta Biodesign #10214) with 30-fold molar excess in HEPES buffer (pH7.0) at 37°C for 1 hour, followed by the removal of the unconjugated linker through 70% ethanol precipitation and size exclusion chromatography. Reduced antibody and modified DNA were combined with the molar ratio of 1:10, and incubated at 37°C for 1 hour and 4°C overnight. The next day, DNA-protein conjugates were purified using protein G affinity chromatography (Genscript #L00209) to remove unconjugated DNA.

Anti-PD-L1 (Bio X cell #BE0285) antibody was conjugated with 3’NH2-5’Quasar570 modified DNA 22mer through a HyNic-4FB linker (TriLink, #S-9011) following the kit instructions. As a validation of the antibody activity post the conjugation, purified anti-PD-L1-DNA conjugates from the “TCEP” strategy and the TriLink strategy were incubated with PD-L1 overexpressing K562 cells at 4°C for 30 minutes, followed by two washes (centrifugation at 400 g x 5 minutes and resuspension in PBS) for flow cytometry with a BD LSR II. Anti-PD-L1-DNA conjugate linked through the “TCEP” strategy was loaded on DNA-scaffolded PLGA microparticles. PD-L1 overexpressed and wildtype K562 cells at 1 million/mL were respectively added with particles at 0.3 OD550 and incubated for 30 minutes at 37°C, followed by cell nuclei staining with Hoechst 33342 (ThermoFisher #62249) and the imaging using the spinning disk confocal fluorescence microscope (Nikon Yokogawa CSU-22).

His-tag GFP were expressed by Escherichia coli BL21 (DE3) (Novagen) transduced with pRSET-EmGFP vector (ThermoFisher, #V35320) in E. coli expression medium (MagicMedia, Invitrogen #K6803), and extracted using cell lysis reagent (Sigma, #B7435) followed by the purification using nickel-nitrilotriacetic acid affinity chromatography (Invitrogen #R90115). The MAL-PEG4-NHS linker (Quanta Biodesign) was mixed with GFP at 30-fold molar excess, and incubated at 37°C for 1 hour followed by the size-exclusion chromatography to remove the excess. 3’Thiol-modified complementary DNA was decapped using the protocol described above, and reacted with modified GFP at 1:10 molar ratio at 37°C for 1 hour followed by 4°C overnight. The next day, GFP-DNA conjugates are purified using nickel-nitrilotriacetic acid affinity chromatography to remove unconjugated DNA. Protein-DNA conjugates were analyzed through SDS-PAGE gel electrophoresis (Genscript #M42012L) with SYPRO Ruby dye staining (ThermoFisher #S21900).

### Ratiometric control of surface DNA scaffolds with different sequences and the co-loading of versatile payloads

3’Thiolated DNA with different sequences were synthesized, namely R, G, and B (**Fig. Supplementary 2f**). Following the conjugation with PLGA10k-PEG5k-Mal (Akina, #AI053), polymer-DNA molecules from different sequences were mixed at varying ratios and incorporated into the particle fabrication process described above (100 nmol for each reactant). Complementary strands to the DNA scaffolds that were pre-conjugated with small molecules (e.g. fluorescent dye or biotin, at 1 μM/OD550) or proteins (e.g. GFP or antibodies, at 180 nM/OD550), were incubated with particles at 37°C for 30 minutes in PBS buffer with 600 mM Na+ and 0.01% Tween-20 supplemented for the surface hybridization of payloads. The excess was removed by three washes. For the co-loading of multiple cargos, the input proportion of each individual one with a specific sequence was consistent with its DNA-polymer counterpart input during the particle fabrication.

For the loading of biotinylated biomolecules on 3’-biotinylated complementary DNA hybridized particles, a large excess of streptavidin (Prozyme #SA10) was added at 1.1 mg/mL per OD550 and incubated at room temperature for 30 mins, followed by three washes. Biotinylated antibody, protein or peptide was added at 180 nM/OD550 and incubated at room temperature for 30 mins to bind with surface streptavidin followed by three washes.

Loading efficiencies of different cargos on-surface or in-core were quantified through the fluorescence-based assay of dye-labeled DNA strands (5’Quasar570-compR, 5’Quasar705-compG, 5’Quasar670-compB, **Fig. Supplementary 2f**), peptides (C-terminus FITC-labeled peptide, LifeTein, LLC) or proteins (FITC-labeled human IgG, Sigma #SLBW7799), after etching of particles by dimethyl sulfoxide (DMSO) and the dilution with 9x volume of water. The fluorescence signal was detected by a plate reader (Tecan Spark) and analyzed using a calibration curve with the normalization from OD550. Functionalized microparticles with fluorescent labels were imaged through a spinning disk confocal fluorescence microscope (Nikon Yokogawa CSU-22), and the size distribution was quantified using ImageJ software analysis of images acquired. Fluorescent microparticles were also analyzed the concentration using a cell counter (Countess II FL AMQAF1000, ThermoFisher).

### Surface step-by-step conjugation for payload attaching

DNA-scaffolded particles were hybridized with 3’amine-modified complementary DNA strands at 1 μM/OD550 followed by three washes. A large excess of MAL-dPEG4-NHS linker (Quanta Biodesign) was added at 30 μM/OD550 and incubated at room temperature for 1 hour followed by three washes to bring maleimide functional groups onto DNA scaffolds. To fabricate the control particles with maleimide functional groups but without DNA scaffolds, 100 nmol block-co-polymer PLGA-PEG-MAL (Akina #AI053) was used to replace the polymer-DNA reaction mixture during the PLGA microparticle fabrication described above. Freshly prepared particles with maleimide functional groups were then reacted with 3’thiol-5’Quasar670-modified DNA at 1 μM/OD550 or reduced antibody with free thiols (FITC-IgG, Sigma) at 180 nM/OD550 for 1 hour at 37°C and 4 °C overnight. The next day particles were washed and quantified through the fluorescence-based analysis.

### DNA scaffolds stability test in DNase and human serum

PLGA microparticles were hybridized with fluorescently labeled complementary DNA strands, followed by human FITC-IgG (Sigma # SLBW7799) loading through surface step-by-step conjugation. Particles with and without IgG coverage were suspended in the enzyme reaction buffer (Promega #M6101) plus DNase (RQ1 RNase-free DNase, Promega #M6101) at 5 U per 1 OD550 × 50 μL and incubated at 37°C for 20 mins, followed by the addition of stop buffer (Promega #M6101) and the centrifugation at 10,000 x g for 10 mins to collect the supernatant and pellet for the fluorescence-based analysis. Similarly, particles with and without IgG attachment together with those hybridized by GFP-DNA conjugates were suspended in human serum (bioWORLD #v13081400) and incubated for 1 hour at 37°C, followed by the centrifugation to collect the supernatant and pellet for the fluorescence-based analysis.

### Cell lines and culture

K562 human myelogenous leukemia cells (ATCC #CCL-243) and A375 human malignant melanoma cells (ATCC #CRL-1619) were used in T cell killing experiments. K562 and A375 were lentivirally transduced to stably express human HER2, PD-L1 and GFP. All cell lines were FACS-sorted for expression of the transgenes. A murine macrophage cell line J77A4.1 (ATCC TIB-67) was used for particle uptake assay. K562 cells were cultured in Iscove’s Modified Dulbecco’s Medium (Corning #10-016-CV) with 10% fetal bovine serum (FBS, UCSF Cell Culture Facility) and gentamicin (UCSF Cell Culture Facility). A375, J77A4.1 cells and Lenti-X 293T packaging cells (Clontech #11131D) were cultured in Dulbecco’s Modified Eagle’s Medium (Gibco #10569-010) with 10% FBS.

### Macrophage uptake assay

Biotin-modified “self”-peptide (biotin-miniPEG-GNYTCEVTELTREGETIIELK[Lys(FITC)], LifeTein) was loaded onto streptavidin-coated PLGA microparticles using the protocol described above. The streptavidin of the control particles without “self-peptide” were labeled with NHS-fluorescein (ThermoFisher #46409) at 6 μM/OD550 for 1 hour at room temperature, followed by three washes. Murine macrophage cell line J77A4.1 (ATCC) grown on glass bottom chamber slides (Thermo Scientific #154526) was treated with lipopolysaccharides (Sigma #L4391) at 100 ng/mL for overnight. On the next day, “self”-peptide loaded particles and control particles were added at 0.03 OD550 × 200 μL and incubated at 37°C for 1 hour. Cells were then washed three times with PBS to remove un-internalized particles and fixed using 4% paraformaldehyde (Electron Microscopy Sciences #15710) for 20 minutes. After three washes, cells were imaged using the spinning disk confocal fluorescence microscope (Nikon Yokogawa CSU-22). Images were analyzed for cell fluorescence signal using ImageJ software.

### Primary human T cell isolation and culture Primary

CD4+ and CD8+ T cells were isolated from anonymous donor blood after apheresis by negative selection (STEMCELL Technologies #15062 and #15063). T cells were cryopreserved in RPMI-1640 (Corning #10-040-CV) with 20% human AB serum (Valley Biomedical, #HP1022) and 10% DMSO. After thawing, T cells were cultured in human T cell medium consisting of X-VIVO 15 (Lonza #04-418Q), 5% Human AB serum, and 10 mM neutralized N-acetyl L-Cysteine (Sigma-Aldrich #A9165) supplemented with 30 units/mL IL-2 (NCI BRB Preclinical Repository) for all experiments.

### Antibody staining and flow cytometry analysis

All antibody staining for flow cytometry was carried out in wells of round bottom 96-well tissue culture plates. Cells were pelleted by centrifugation of plates for 4 minutes at 400 x g. Supernatant was removed and cells were resuspended in 50 μL PBS containing the fluorescent antibody of interest. Cells stained 25 minutes at 4°C in the dark. Stained cells were then washed two times with PBS and resuspended in fresh PBS for flow cytometry with a BD LSR II. All flow cytometry data analysis was performed in FlowJo software (TreeStar).

### AICE-mediated activation of human primary T cells for *ex vivo* expansion

AICEs were synthesized with PLGA microparticles functionalized with anti-CD3 antibody (Bio X Cell #BE0001-2, attached to 3’NH_2_-5’Qusar670-modified compB), and anti-CD28 antibody (Bio X Cell #BE0248, attached to 3’NH_2_-5’FAM-modified compR) at varying ratios from 1:5, 1:3, 1:1, 3:1 to 5:1 according to the method described above. After thawing and 24-hour recovery in culture, 1.4 × 10^5^ CD4+/CD8+ human primary T cells in 200 μL medium were added with AICEs at 0.11 OD550 (approximately 2.5 AICE to 1 cell), or CD3/CD28 Dynabeads at a 1:2.5 cell:bead ratio (Life Technologies #11131D) for 4 days. CD8+ T Cells were imaged through the spinning disk confocal microscope (Nikon Yokogawa CSU-22) one day after AICE addition. CD4+ and CD8+ T Cells numbers were quantified using a cell counter (Countess II FL AMQAF1000, ThermoFisher) every other day from day 6 to day 14 after AICE activation. Fresh IL-2 containing medium was supplemented to maintain the cell concentration between 0.5-1.5 million per mL. On day 14, T cell phenotype was studied by flow cytometry using the following antibodies: anti-CCR7 (BD #561271), anti-CD45RA (BD #562885), anti-LAG-3 (BD #565720), anti-TIM-3 (BD #565558), anti-PD-1 (Biolegend #329936).

Anti-IL-2 clone #5355 (ThermoFisher #MA523696) was biotinylated using NHS-PEG_4_-biotin (Quanta Biodesign #10200) at 30 x molar excess, with the incubation at 37°C for 1 hour followed by size exclusion chromatography (ThermoFisher #89882) to purify. Streptavidin-coated PLGA microparticles were incubated with biotinylated anti-IL-2 at 20 nM/OD550 and IL-2 (NCI BRB Preclinical Repository) at 30 nM/OD550 at room temperature for 30 minutes, and washed twice. Surface-bound IL-2 on particles were supplemented at 0.1 OD550/mL to AICE [3:1]-treated T cells and replenished to maintain the cell concentration at 0.5-1.5 million per mL, as a comparison to free IL-2 supplementation at 30 units/mL.

### Transduction of synthetic Notch CAR T cells

The binding domains of LaG17 nanobody^28^ and GCN4 (PNE) scFv^42^ were cloned into synNotch receptors with Gal4 DNA-binding domain VP64 as a synthetic transcription factor^28^. The response element plasmids was modified from the pHR’SIN:CSW vector with five copies of the Gal4 DNA binding domain, a HER2-CAR in the multiple cloning site downstream of the Gal4 response elements, and a PGK promoter that constitutively drives mCherry expression to easily identify transduced T cells^28^. All constructs were cloned via InFusion cloning (Takara Bio #638910). Pantropic VSV-G pseudotyped lentivirus was produced via transfection of Lenti-X 293T cells with a pHR’SIN:CSW transgene expression vector and the viral packaging plasmids pCMVdR8.91 and pMD2.G using Fugene HD (Promega #E2312). Primary T cells were thawed the same day and, after 24 hours in culture, were stimulated with Human T-Activator CD3/CD28 Dynabeads (Life Technologies #11131D) at a 1:2.5 cell:bead ratio. At 48 hours, viral supernatant was harvested and the primary T cells were exposed to the virus for 24 hours. At day 5 after T cell stimulation, Dynabeads were removed and the T cells expanded until day 12 when they were rested and could be used in assays. T cells were sorted for assays with a FACs ARIA II on day 5 post T cell stimulation.

### *In vitro* stimulation of synNotch T cells through AICEs

2.5 × 10^4^ synNotch CD4+ or CD8+ T cells were co-cultured with target cancer cells at a 1:1 ratio, with the addition of AICE at 0.075 OD550 × 200 μL medium (100 μL T cell medium + 100 μL cancer cell medium) (unless otherwise specified). Dual antigen (GFP and HER2) positive target cells (A375 and K562) were used as positive controls. After mixing, cells were centrifuged for 2 min at 300 x g to initiate interaction of the cells. After 24 hours, the co-culture from CD8+ T cells were stained with anti-CD69 antibody (BD #562884) and analyzed by flow cytometry (BD LSR II), and also imaged through the spinning disk confocal microscope (Nikon Yokogawa CSU-22). After 48 hours, cytokine concentration in the supernatant was measured by IL-2 ELISA (for CD4+ T cells, eBiosciences #BMS2221HS) and Interferon-γ (IFN-γ) ELISA (for CD8+ T cells, ThermoFisher #KHC4021).

To quantify target cell killing, 2.5 × 10^4^ A375 cells were seeded on flat-bottom 96-well tissue culture plate (Falcon #353072) and cultured for 8 hours, followed by the addition of 2.5 × 10^4^ CD8+ T cells and AICE at 0.075 OD550 × 200 μL final (unless otherwise specified). After 1-3 days, cells were gently washed with PBS for 2 times, and analyzed for cell viability using PrestoBlue Cell Viability Reagent (Invitrogen #A13262).

### IVIS imaging of IR dye-labeled AICE in mice tumors

NH_2_-modified PLGA polymer (LG 50:50, Mw 30,000-40,000 Da) dissolved in DMF was reacted with IR800CW-NHS Ester (Li-COR #P/N 929-70020) at 30-fold molar excess at room temperature for 1 hour, followed by repeat 70%-ethanol precipitation and DMF re-dissolving to remove the unconjugated dye. 1 mg of IR800CW-labeled polymer was incorporated in the emulsion protocol for DNA-scaffolded (G strand) PLGA microparticles, and 5’Quasar705-modified compG strand was hybridized on surface. NSG mice (Jackson Laboratory #005557, female, ~8-12-weeks old) were implanted with xenograft tumors - 5 × 10^6^ K562 tumor cells subcutaneously on the left and right flank. 10 days after tumor implantation, 50 μL particles at 100 OD550 were injected intratumorally, and imaged under IVIS 100 preclinical imaging system (Xenogen #124262) every 3-4 hours for the first 48 hours and every 8 hours for the rest of the week. Images are analyzed using Living Image Software (PerkinElmer).

### *In vivo* tumor targeting

NSG mice (Jackson Laboratory #005557, female, ~8-12-weeks old) were implanted with two identical xenograft tumors - 5 × 10^6^ HER2+ K562 tumor cells subcutaneously on the left and right flank. Seven days after tumor implantation, human primary CD4+ (4 × 10^6^) and CD8+ T cells (4 × 10^6^) were injected intravenously into the tail vein of the mice. These T cells were either untransduced (control) or engineered with the anti-GFP synNotch Gal4VP64 receptor and the corresponding response elements regulating anti-HER2 4-1BBζ CAR expression. On the same day, AICE-GFP particles were injected intratumorally at one side of the two flanks with 50 (or 10) OD550 × 50 μL per dose, leaving the other as the control. Three additional doses of AICE were injected into the same tumor every 4 days or starting as the tumor grew over 500 mm^3^ in volume. Tumor size was monitored via caliper over 20 days after T cell injection. Mice were euthanized as any tumor reached 20 mm in diameter or became ulcerated.

Remaining tumors from contralateral flank and early euthanized mice were collected and fixed in 4% paraformaldehyde (Electron Microscopy Sciences #15710) overnight, followed by the dehydration in 30% sucrose (Sigma #S9378) for 1-2 days until sunk. Tumors were then snap-frozen in OCT (Tissue-Tek) to produce cryosections. Sections were stained with anti-CD3 antibody (Biolegend #344811) in 1% BSA/PBS for 1 hour and washed three time, followed by the imaging with the spinning disk confocal fluorescence microscope (Nikon Yokogawa CSU-22).

### Statistical analysis

Statistical analysis were preformed using GraphPad Prism software. All values and error bars are mean ± s.d., except where indicated differently. Two-tailed paired *t* test, one-way ANOVA test, one site-specific binding model fit, and fourth order polynomial model fit were performed where appropriate. Life Sciences Reporting summary. Further information on experimental design is available in the Nature Research Summary linked to this article.

### Data availability

The authors declare that the data supporting the findings of this study are available within the paper (and its supplementary information files).

