## Supplemental information for "DNA-scaffolded biomaterials enable modular and tunable control of cell-based cancer immunotherapies"

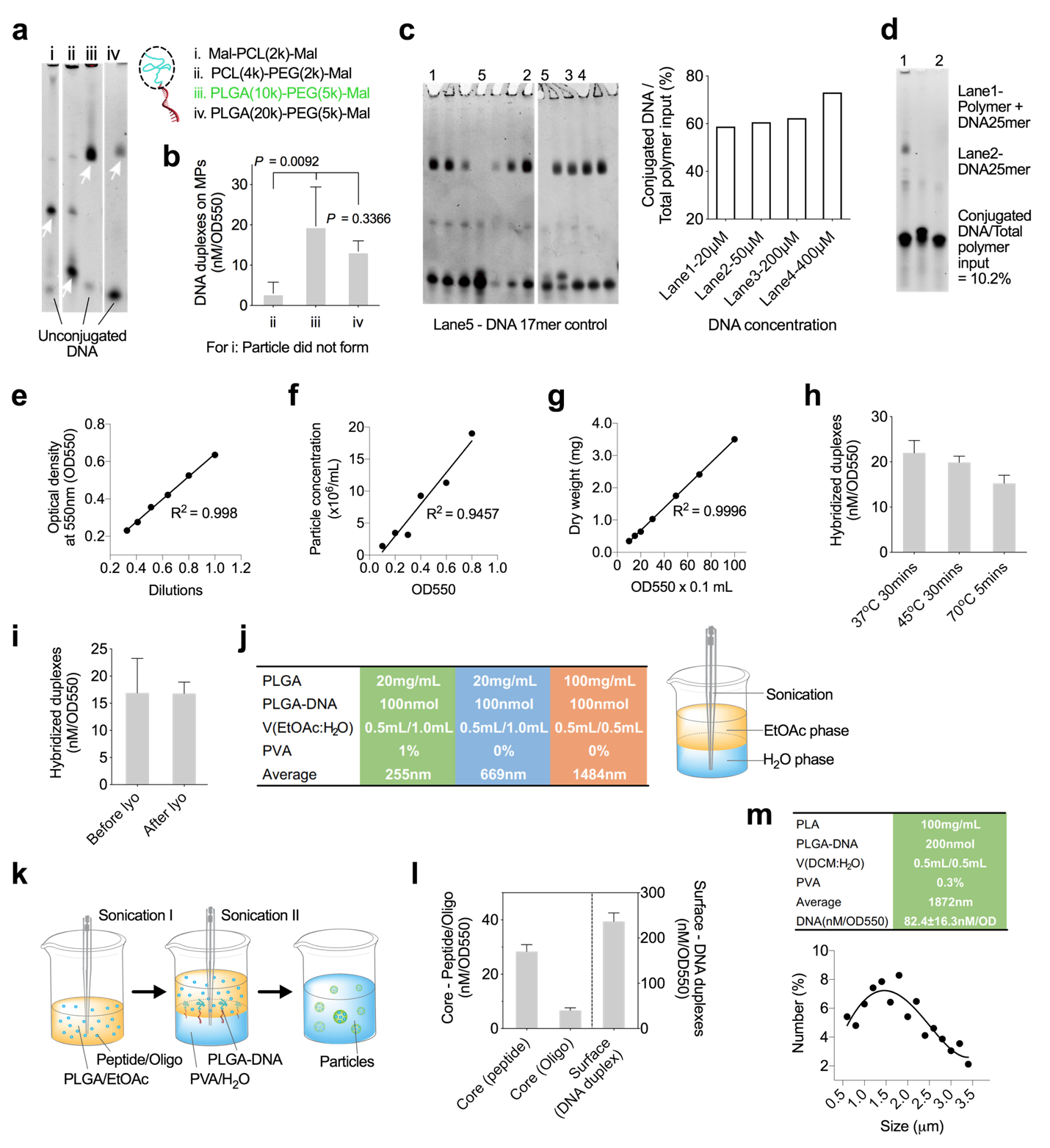

Supplementary Figure 1 Optimization of DNA scaffolds on polymeric particles

(**a**) Representative denatured TBE-Urea PAGE gel image of different polymer-DNA conjugates yielded from the conjugation of maleimide(Mal)-modified polymer at 100 μM with thiol-modified DNA 17mer at 20 μM (n = 3 independent experiments). White arrows, product bands. Modified polymers tested are Mal-PCL(polycaprolactone, Mw 2,000)-Mal^6^, PCL(Mw 4,000)-PEG(polyethylene glycol, Mw 2,000)-Mal (Akina #AI082), PLGA(poly(lactic-co-glycolic acid), 50:50, Mw 10,000)-PEG(Mw 5,000)-Mal (Akina #AI053) and PLGA(Mw 20,000)-PEG(Mw 5,000)-Mal (NSP #12191). (**b**) Hybridized surface DNA on PLGA microparticles (target 1-2 μm in diameter) by incorporation of different polymer linkers for particle fabrication. Data are mean ± s.d. (n = 3 independent experiments). The polymer-DNA conjugate made from Mal-PCL(2k)-Mal did not yield particles at the desired size, while for PCL(4k)-PEG(2k)-Mal, 0.2% PVA (polyvinyl alcohol) was required during the fabrication. (**c-d**) Denatured TBE-Urea PAGE gel image of polymer-DNA reaction (DNA:polymer=1:1) at varying reactant concentrations (**c**) and 25mer DNA (**d**), quantified by densitometry analysis (n = 1 for each variable). Reactant at 400μM formed obvious aggregates, which may over-estimate the conjugation efficiency, and 25 mer yielded lower conjugation efficiency than 17 mer (**Fig. 1c**). Thus, reactants at 200 μM and DNA-17mer was selected for later reactions. (**e-g**) Linear regression of the dilution (**e**), particle concentration by Cell Counter (ThermoFisher) (**f**) and dry weight (**g**) of PLGA microparticles to optical density at 550 nm (OD550). n = 1 for each variable in (**c-g**). (**h-i**) Hybridized DNA scaffold on PLGA microparticles from different hybridization conditions (**h**) and lyophilization (lyo) treatment (**i**). Data are mean ± s.d. (n = 3 independent samples). (**j**) Schematic of the size control of DNA-decorated PLGA particles by varying key parameters in the fabrication protocol. The size can be tuned through the varying of polymer concentration, the two-phase volume and ratio, and the employment of other surfactants (e.g. PVA). (**k**) Schematic of the double-emulsion protocol to incorporate biomolecules in the core of DNA-scaffolded particles, where small volume of peptides and oligonucleotides (oligo) are emulsified in the organic solvent prior to the polymer-DNA emulsion. (**l**) Fluorescence quantification of dye-labeled peptide and oligo in the core, compared to surface-hybridized DNA scaffolds. Data are mean ± s.d. (n = 5 from two independent experiments). (**m**) The emulsion protocol for DNA-scaffolded microparticles made of polylactic acid (PLA), and their size distribution by ImageJ software analysis of confocal fluorescence images of particles hybridized with dye-labeled complementary DNA strands (n = 6 technical replicates of 3 independent samples). The curve is a non-linear fit by fourth order polynomial model.

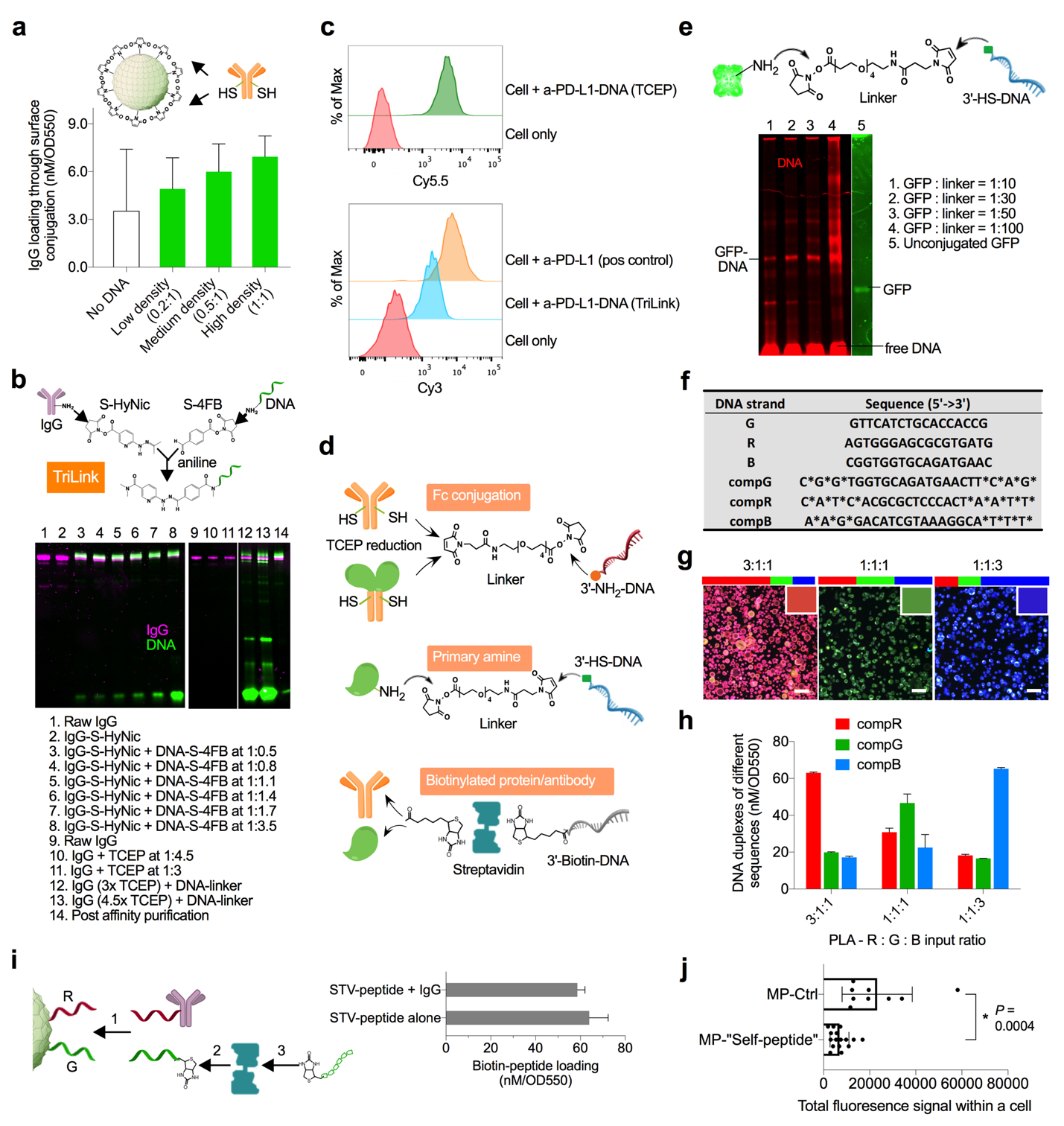

Supplementary Figure 2 Surface functionalization strategies for various protein surface conjugates

(**a**) Quantification of the surface conjugation of FITC-labeled IgG to the functional groups present on PLGA microparticles with and without DNA scaffold at different densities. Data are mean ± s.d. (n = 3 independent experiments). (**b**) SDS-PAGE gel image of DNA-IgG (anti-PD-L1) conjugates linked through different chemistries: TriLink (to 3’NH_2_-5’Quasar570 DNA 22mer) versus “TCEP” (to 3’NH_2_-5’Quasar705 DNA 22mer). Optimal DNA to IgG ratio was essential for the protein activity through the TriLink strategy, and protein aggregation was observed at higher DNA to IgG ratio. TCEP strategy led to some IgG fractionation, but can be minimized through lowering TCEP to IgG ratio. Unattached DNA were removed by affinity chromatography purification prior to the surface hybridization. Data represent n = 3 independent experiments. (**c**) Flow cytometric analysis of PD-L1 overexpressing K562 stained with DNA-antibody (anti-PD-L1) linked by the TriLink and “TCEP” strategies, in comparison to commercial PE-labeled anti-PD-L1 (n = 1 for each variable). (**d**) Conjugation strategies of complementary DNA to proteins and antibodies. The hinge region of IgG or Fc-tagged proteins can be partially reduced to expose thiols for tethering with amine-modified DNA through the Mal-PEG_4_-NHS linker; proteins can be linked through primary amines to thiol-modified DNA via the Mal-PEG_4_-NHS linker; biotinylated proteins or antibodies can be attached on particle surface through the streptavidin bridge. The “TCEP” strategy was extensively used in this work for many antibodies to generate DNA-protein conjugates without activity loss. In terms of therapeutic proteins other than antibodies, a commonly found Fc-tag on commercial proteins can be utilized with the same chemistry; for certain protein sensitive to TCEP reduction, an antibody to its Fc-tag or non-essential site can be first incorporated as the surface dock; otherwise, proteins can also be modified on their primary amines, but the excess ratio of the linker needs to be optimized to maintain the activity. (**e**) SDS-PAGE gel image of GFP-DNA conjugates synthesized through the linking of primary amine to thiol-modified DNA with different linker excess. Data represent n = 2 independent experiments. (**f**) Sequence of DNA strands (G, R and B) that are attached to polymers, and their respective complementary strands (compG, compR and compB). *: phosphorothioated DNA base. (**g**) Confocal fluorescence microscopy images of PLA microparticles with DNA scaffolds of different sequence compositions post hybridization with the corresponding dye-labeled compDNA. The merged images of particles agreed well with the theoretically integrated color (upper right) at different input ratios. (**h**) DNA duplexes of different sequences on PLA particles by fluorescence-based quantification. Data are mean ± s.d. (**g** and **h** are from n = 2 independent experiments). (**i**) Fluorescence-based quantification of biotinylated peptide loaded through streptavidin on DNA scaffolds (G), with and without the co-loading of IgG on DNA scaffolds of a different sequence (R). Data are mean ± s.d. (n = 3 independent samples). (**j**) Fluorescence-based quantification of particles in individual J774A.1 macrophage cells through ImageJ software analysis. Data are mean ± s.d. (n = 3 biologically independent samples per group) and *P* value was determined by two-tailed paired *t* test.

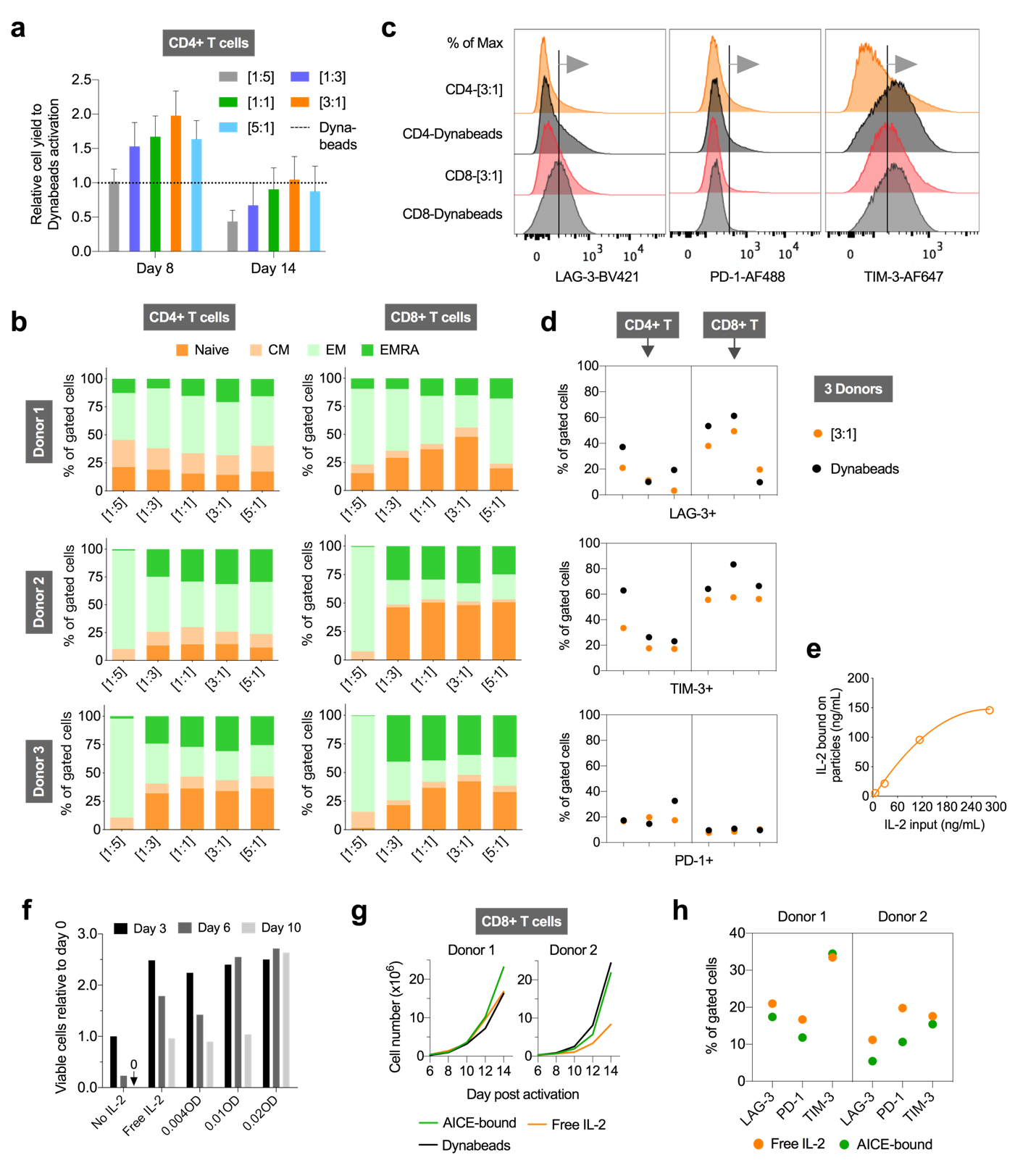

Supplementary Figure 3 Activation of human primary T lymphocytes by functionalized AICE for *ex vivo* expansion.

(**a**) Cell yield of CD4+ T cells at day 8 and 14 after activation by AICE with CD3 and CD28 antibodies at different ratios, relative to Dynabeads. Data are mean ± s.e.m. (n = 3 independent donors of two independent experiments). (**b**) Differentiation profile of T cells from 3 individual donors at 14 days after activation by AICE with varying ratios of surface stimulatory moieties. (**c**) Gating strategy of the exhausted cell population based on expression levels of LAG-3, PD-1 and TIM-3 on expanded CD4+/CD8+ cells 14 days after activation by AICE [3:1] and Dynabeads. (**d**) Exhaustion phenotype of CD4+/CD8+ T cells from 3 individual donors 14 days after activation by AICE [3:1] and Dynabeads. (**e**) Binding curve of IL-2 to PLGA microparticles coated with IL-2 antibodies. Data are mean ± s.d. (n = 4 technical replicates from 2 independent samples), and the curve is a nonlinear fit by the one site-specific binding model. (**f**) Viable CD8+ human primary T cells over time with different doses of surface bound IL-2 on PLGA microparticles versus free IL-2. The amount of free IL-2 is equivalent to particles between 0.004 OD and 0.01 OD. Data are mean of n = 2 technical replicates. (**g**) Growth curve of CD8+ primary T cells from two different donors treated with AICE [3:1] together with AICE-bound IL-2 or free IL-2 at equivalent dose, compared to Dynabeads with free IL-2 control. Data are mean of n = 2 technical replicates. (**h**) Levels of LAG-3, PD-1 and TIM-3 expression for CD4+ T cells of two different donors 14 days after activation by AICE [3:1] together with AICE-bound IL-2, or free IL-2.

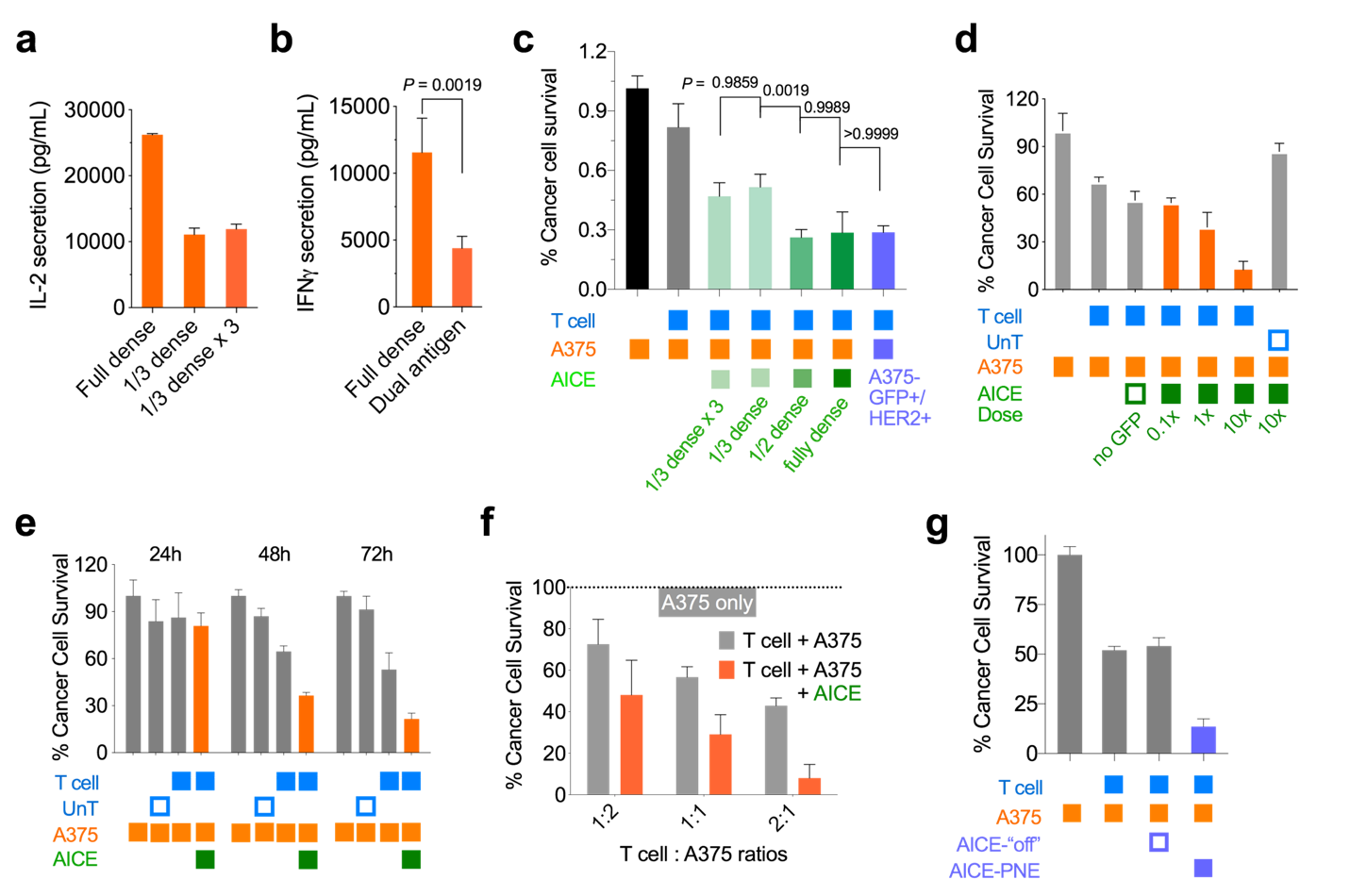

Supplementary Figure 4 Activation of human primary synNotch CAR-T cells for AND-gated target cell toxicity using AICE coated with priming antigens.

(**a**) IL-2 secretion levels by primary synNotch CAR CD4+ T cells 48 hours post co-incubation of A375-HER2 and AICE with different GFP densities and different particle doses. Total amount of antigen presented by AICE particles with “Fully dense” was equal to “1/3 dense x 3”. The antigen density on AICE matters more than the total amount of antigen for synNotch CAR-T cell activation. Data are mean ± s.d. (n = 2 technical replicates from 4 biologically independent samples). (**b**) IFN-γ secretion concentration from primary CD8+ T cells 48 hours post co-incubation of AICE and A375-HER2 versus dual antigen target cells A375-GFP+/HER2+. Data are mean ± s.d. (n = 4 technical replicates from 4 biologically independent samples), and *P* value was determined by two-tailed paired *t* test. (**c-f**) Cell survival study of target A375-HER2 cells from co-culturing of AICE-primed synNotch CAR-T cells, with a series of conditions, including varying densities of priming antigen on AICE (**c**), different doses of AICE to T cells (**d**), different duration of treatments (**e**) and varying T cell to target cell ratios (**f**). Data are mean ± s.d. (n = 4 biologically independent samples). (**g**) Cell survival test of target A375-HER2 cells treated with a different type of synNotch CAR-T cells responsive to PNE peptide present on AICE after 48 hours of co-culture. Data are mean ± s.d. (n = 4 biologically independent samples).

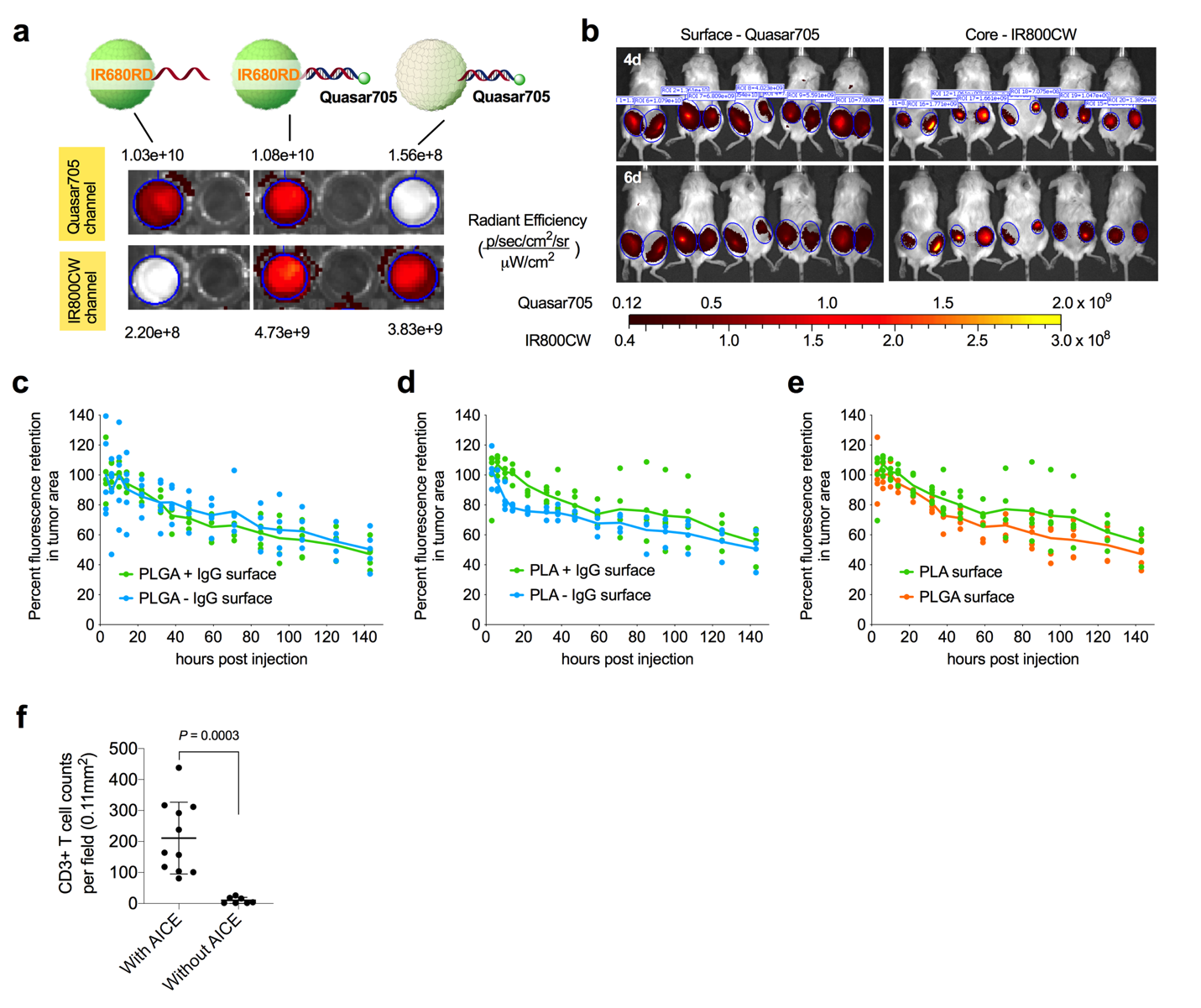

Supplementary Figure 5 *In vivo* half-life of AICE with different compositions and their priming of synNotch CAR-T cells for selective tumor clearance.

(**a**) IVIS near-infrared fluorescence imaging of AICE *in vitro* with Quasar705 on surface and IR800CW in core. The two dyes can be imaged at the same time without cross-interference. (**b**) IVIS fluorescence imaging of labeled AICE in subcutaneous K562 tumors of NSG mice at day 4 and day 6 (n = 5 mice). Left tumors are injected with AICE made of PLA; right tumors are injected with AICE made of PLGA. (**c-e**) Quantification of the average fluorescence signal of surface coatings at the tumor area over 6 days, from the injection of PLGA microparticles with and without IgG coverage (**c**), PLA microparticles with and without IgG coverage (**d**), and microparticles made of PLGA versus PLA (**e**). Half-life of the surface coatings of AICE in K562 tumors of NSG mice was not affected by the polymer composition (between PLGA and PLA) and surface IgG coverage. Lines are the mean signal from n = 5 mice. (**f**) Quantification of CD3+ T cell numbers in the stained tumor sections from mouse 8 shown in Fig. 5h (n = 10 technical replicates).
